# *dArc1* controls sugar reward valuation in *Drosophila melanogaster*

**DOI:** 10.1101/2024.11.04.621761

**Authors:** Sven Bervoets, Miles Solomon Jacob, Anita V. Devineni, Brennan Dale Mahoney, Kaelan R. Sullivan, Andrew R. Butts, Hayeon Sung, Jenifer Einstein, Mark M. Metzstein, Monica Dus, Jason D. Shepherd, Sophie Jeanne Cécile Caron

## Abstract

The *Arc* genes — which include *Drosophila Arc1* and *Arc2* (*dArc*) — evolved from Ty3 retrotransposons and encode proteins that form virus-like capsids. These capsids enable a novel form of intercellular communication by transferring RNAs between cells. However, the specific neuronal circuits and brain processes Arc intercellular signaling regulates remain unknown. Here, we show that loss of both *dArc* genes in *Drosophila melanogaster* enhances associative learning in an appetitive conditioning paradigm, where flies associate an odor with sugar rewards. This increased learning performance arises from an increased valuation of sugar rewards: unlike wild-type flies, *dArc^-/-^* flies form abnormally strong associations even when the sugar reward is small or has no caloric value. We found that the γ5-dopaminergic neurons of the protocerebral anterior medial (PAM) cluster, which encode the positive valence of sugar rewards, show heightened activity in response to sucrose in *dArc^-/-^* flies. We further show that the learning phenotype of *dArc^-/-^* flies depends on the formation of capsids, underscoring a direct role for capsid-mediated Arc signaling in sugar valuation. Our findings establish *dArc* genes as critical regulators of reward valuation in *D. melanogaster*, acting through a non-cell autonomous mechanism that relies on capsid-mediated communication between cells.

## MAIN TEXT

The activity-regulated cytoskeletal-associated (*Arc*) gene was initially identified in mammals as an immediate early gene — rapidly induced by neuronal activity — that has critical roles in synaptic plasticity^1,2^. Two Arc genes, *dArc1* and *dArc2,* have been identified in schizophoran flies, including *Drosophila melanogaster*. Recent evidence indicates that Arc proteins can self-assemble into capsids, enabling the transfer of RNAs between neurons^3,4^. This non-synaptic signaling mechanism may represent a completely new mode of intercellular communication in nervous systems^5^. The ability to form capsids that can transfer RNAs between cells is an ancestral property of these proteins^4,6^. *Arc* genes evolved from Ty3 retrotransposons — a class of mobile genetic element — and exhibit high sequence homology to Gag domains, which form the capsids of retrotransposons and retroviruses^3,7,8^. However, it remains unclear whether capsid-mediated signaling is essential for the functions of *Arc* genes.

*dArc1* and *dArc2* play diverse roles, influencing processes ranging from metabolism regulation to associative learning^9–11^. However, only *dArc1* has been linked to synaptic function at the larval neuromuscular junction, and its expression has been shown to increase in response to neuronal activity^12,4^. Structural analyses of dArc capsids have resolved their architecture at atomic resolution, identifying key domains for capsid assembly and RNA binding^6^. In the larval neuromuscular junction, dArc1 has been shown to transfer RNAs, including its own mRNA, from motor neurons to muscle cells, highlighting its capacity for intercellular RNA transport^4^. However, it remains unclear whether and how Arc signaling operates in the *Drosophila* brain. Here, we demonstrate that *dArc1* is required in the adult *Drosophila* brain for accurate valuation of sugar rewards during associative learning, and that this function relies on its ability to form capsids.

### dArc1 is expressed in a subset of serotonergic neurons

To determine where dArc1 is expressed in the adult *Drosophila* brain, we performed immunostaining on the brains of five-day-old flies using custom polyclonal antibodies specific to dArc1 (Extended Data Fig. 1a). In brains immunostained for dArc1, we identified on average 29 ± 1 cells with dArc1 expression (Fig. 1a,b and Extended Data Fig. 1b). The somata of these cells are sparsely distributed in the anterior and posterior brain.

**Figure 1.**
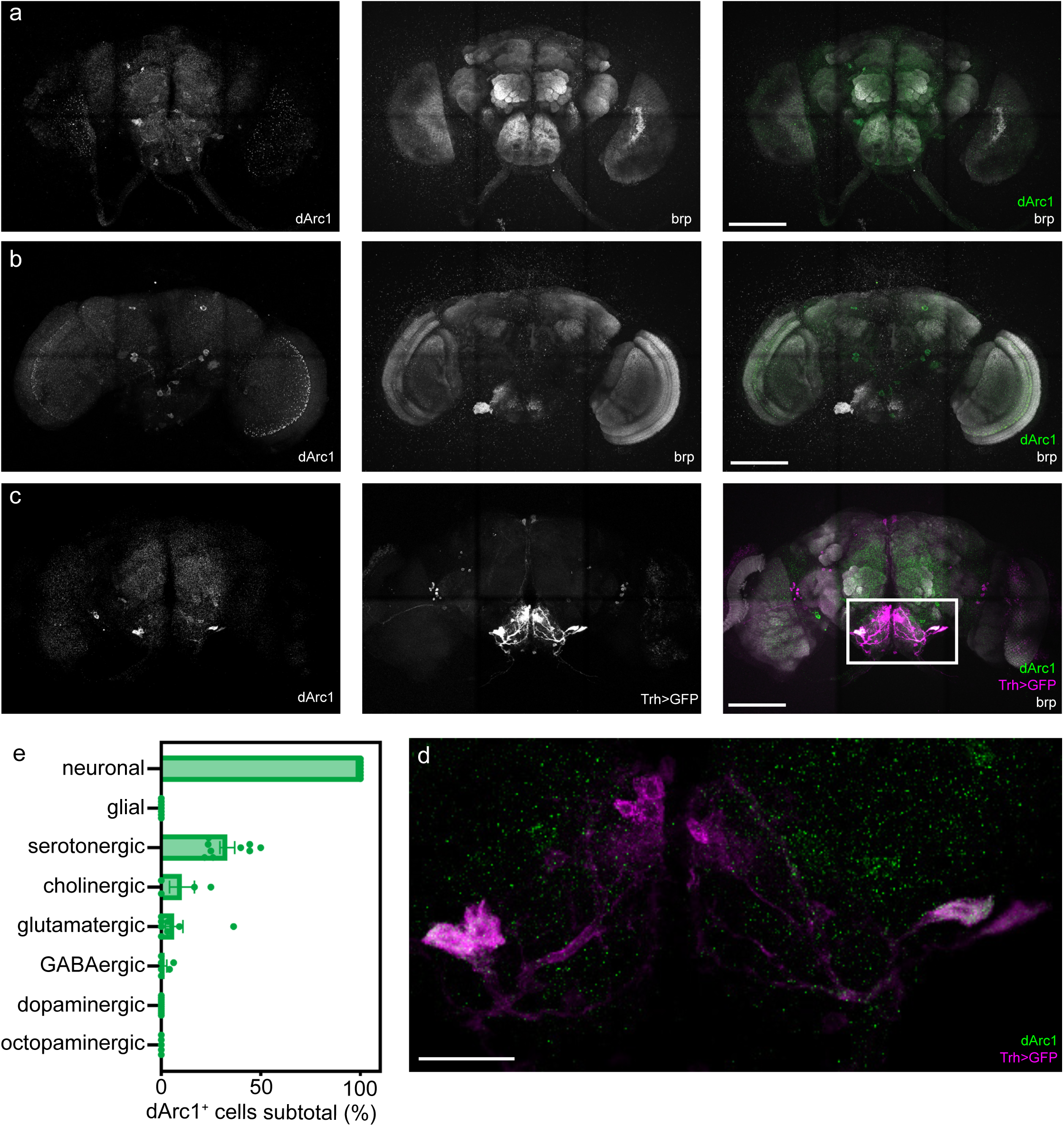
dArc1 is expressed in a subset of serotonergic neurons. (a) The brains of *dArc^+/+^*and *dArc^-/-^* flies were immunostained and imaged using antibodies against the neuropil marker bruchpilot (white) and dArc1 (green) on the anterior and (b) posterior side. Scale bars are 100 µm. (c) The brain of a *Trhn*-GAL4;UAS-GFP fly was immunostained and imaged using antibodies against bruchpilot (white) and dArc1 (green) and intrinsic GFP fluorescence signal (magenta). Scale bar is 100 µm. (d) Zoomed-in image showing the colocalization of dArc1 (green) in the serotonergic neurons (magenta) of the suboesophageal zone. Scale bar is 50 µm. (e) The brains of flies carrying a GAL4 transgene and the UAS-GFP transgene were immuno-stained and imaged using antibodies against dArc1 and intrinsic GFP signal; cells were typed based on whether they express GFP and the fraction of the dArc1-expressing cells belonging to a given type was measured (n ≥ 4 flies). Subjects were mixed sex. Bar charts are presented as means ± s.e.m.

Using cell-type specific reporter transgenes, we found that dArc1 expression was restricted to neurons and not detected in glial cells (Fig. 1e). Immunostaining for the neuronal marker elav and the glial marker repo corroborated these findings (Extended Data Fig. 2a,b). Notably, approximately one-third of dArc1-expressing neurons co-expressed the *Trhn-GAL4* transgene, which labels all serotonergic neurons but not all serotonergic neurons express dArc1 (Fig. 1c-e). Four dArc1-positive serotonergic neurons were consistently observed in the suboesophageal zone, a taste processing center in the *D. melanogaster* brain (Fig. 1c,d). We did not detect dArc1 expression in dopaminergic, GABAergic, or octopaminergic neurons, and only a small subset of cholinergic and glutamatergic neurons showed dArc1 expression (Fig. 1e and Extended Data Fig. 2c,d). Thus Arc1-expressing neurons is a heterogeneous group of neurons that consists of serotonergic, cholinergic, and glutamatergic neurons as well as other unidentified neurons.

### *dArc^-/-^* flies outperform *dArc^+/+^* flies in an appetitive conditioning paradigm

*dArc1* and *dArc2* share high homology, with both encoding proteins capable of forming capsids^6^. To prevent compensatory effects from the deletion of only *dArc1*, we used CRISPR-mediated homologous recombination to generate fly lines in which the entire *dArc* locus, a region containing both *dArc1* and *dArc2*, was deleted (*dArc^-/-^* flies; Extended Data Fig. 3a). We also generated a control line, in which the *dArc* locus was left intact (*dArc^+/+^* flies). qPCR of cDNA, western blots of protein extracts, and immunostaining of brains confirmed the absence of *dArc1* and *dArc2* in *dArc^-/-^* flies (Extended Data Fig. 1a and Extended Data Fig. 3b-d). *dArc^-/-^* flies are viable, develop normally and are fertile. They display no visible morphological defects and have normal motor behavior (Extended Data Fig. 3e).

We used a well-established associative conditioning paradigm to test whether *dArc^-/-^* flies have learning defects^13,14^ (Extended Data Fig. 4a). Flies were trained to associate an odor with a sucrose reward, and their memory was tested either 30 minutes (short-term memory) or 24 hours (long-term memory) after training. *dArc^-/-^* flies significantly outperformed *dArc^+/+^* flies in both when their memories were tested after 30 minutes or 24 hours, showing performance indices flies were almost twice as high as those of *dArc^+/+^*flies (Fig. 2a,b).

**Figure 2.**
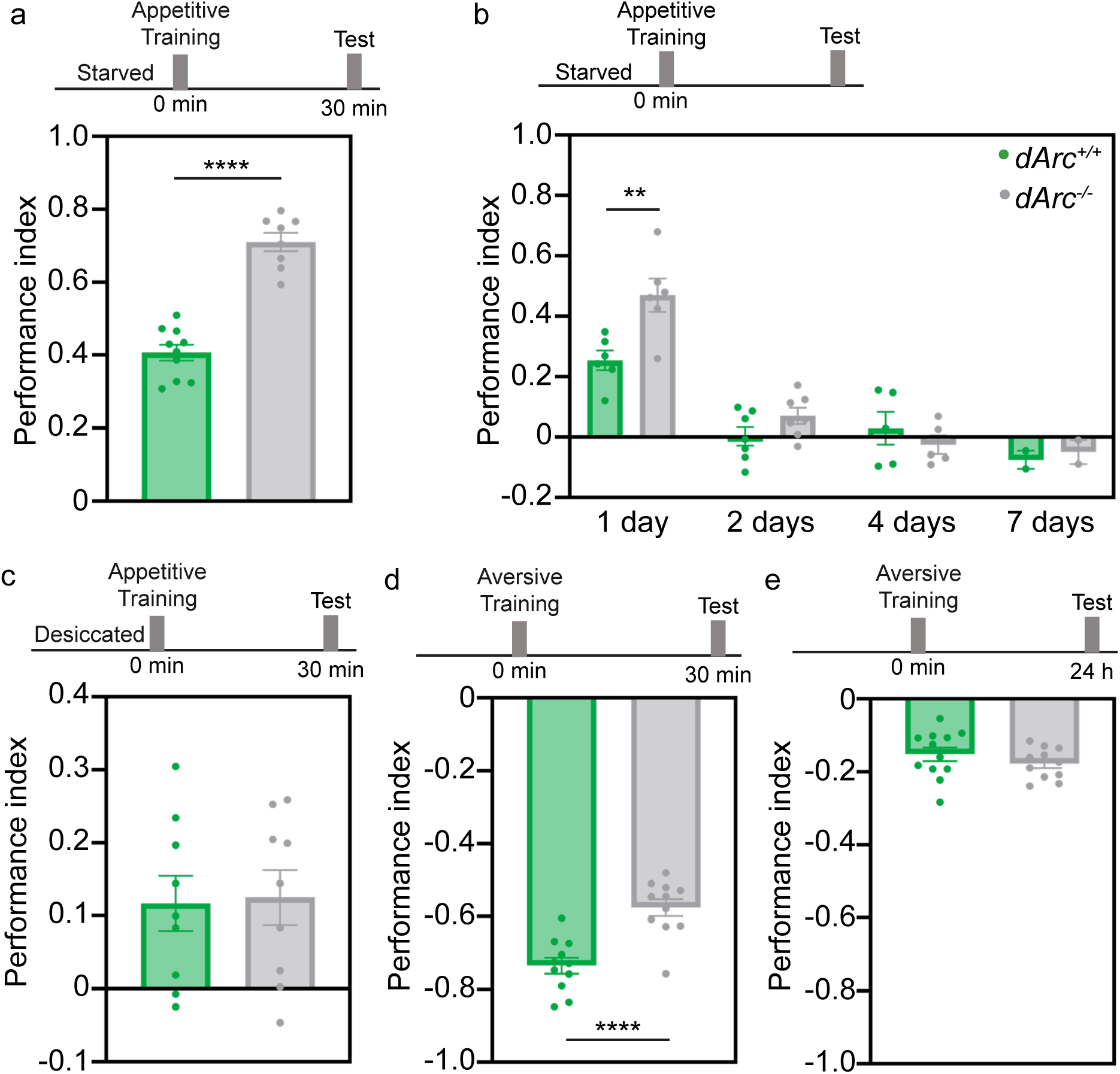
*dArc^-/-^* flies show enhanced sugar reward learning. (a) Flies were starved for 20 hours and subjected to an appetitive association paradigm using sucrose and tested after 30 minutes. (n ≥ 8 groups of flies, Unpaired t-test). (b) Flies were starved for 20 hours and were subjected to an appetitive association paradigm using sucrose and tested after 1, 2, 4, or 7 days. (n ≥ 2 groups of flies, Two-way ANOVA). (c) Flies were desiccated for 20 hours and were subjected to an appetitive association paradigm using water and tested after 30 minutes. (n ≥ 9 groups of flies, Unpaired t-test). (d) Flies were subjected to an aversive association paradigm using a single shock regime and tested after 2 minutes. (n ≥ 10 groups of flies, Unpaired t-test). (e) Flies were subjected to an aversive association paradigm using multiple spaced shock regimes and tested after 24 hours. (n ≥ 11 groups of flies, Unpaired t-test). Subjects were mixed sex. Bar charts are presented as means ± s.e.m.

To determine whether this enhanced learning performance resulted in enhanced memory consolidation, we tested flies two, four, and seven days after training (Fig. 2b). After two days, the appetitive memory in *dArc^+/+^* flies was completely extinguished, while *dArc^-/-^* flies only weakly expressed the memory. However, after four days, the memories in both genotypes were entirely extinguished, suggesting that memories do not persist longer in *dArc^-/-^* flies. To ensure that the difference in performance was not due to variation in sensory perception, we measured odor acuity and innate responses to sucrose and found it to be similar across genotypes and not due to metabolic defects such as abnormal fat mass (Extended Data Fig. 4b-i). Therefore, the enhanced learning of *dArc^-/-^* flies does not result from sensory defects and does not translate into enhanced memory persistence.

To determine whether the enhanced learning phenotype is specific to sucrose rewards or reflects more general defects in reward learning, we tested flies with different reward learning paradigms. We used water as an alternative appetitive reward by modifying the appetitive conditioning paradigm based on previous studies^15,16^. Flies were water-deprived for 20 hours before training, then a water reward was delivered during the training period. Flies were tested 30 minutes after training. We found that *dArc^-/-^*and *dArc^+/+^* flies performed similarly, showing no differences in performance or water preference (Fig. 2c and Extended Data Fig. 5a). This suggests that the defects in *dArc^-/-^* flies might be specific for the sugar reward pathway. To test whether other forms of learning are affected in *dArc^-/-^* flies, we trained flies in an aversive conditioning paradigm using electric shocks^14,17^. We observed a significant reduction in performance indices compared to *dArc^+/+^* flies when we tested short-term memory, but no differences when we tested long-term memory (Fig. 2d,e). Additionally, we observed a significant difference in the ability of *dArc^-/-^* flies to detect electric shocks, a sensory defect that may further contribute to impaired aversive learning (Extended Data Fig. 5b-c).

### *dArc^-/-^* flies do not differ in their hunger and motivational states

The higher sugar reward learning performance indices observed in *dArc^-/-^* flies might result from a dysregulation of the mechanisms assessing nutritional status — *dArc^-/-^* flies might simply be hungrier than *dArc^+/^*^+^flies — or from a dysregulation of the mechanisms that repress appetitive memories when flies are sated. In the first scenario, *dArc^-/-^* flies would constantly be in a heightened hunger state and might express their appetitive memories more readily than *dArc^+/+^* flies, leading to higher performance indices. However, *dArc^-/-^* flies did not show abnormal responses to sucrose: they did not show a greater preference for dwelling over sweet surfaces, nor did they exhibit an aberrant proboscis extension-reflex, suggesting that they were not hungrier than starved *dArc^+/+^* flies (Extended Data Fig. 4b-d). To investigate further possible differences in hunger state, we measured food intake in starved flies that were fed blue-dyed sucrose for two minutes, the same duration as the training phase in the appetitive conditioning paradigm (Extended Data Fig. 4h). We found no significant differences in feeding initiation between the genotypes. Additionally, we detected no significant differences in triglyceride or protein levels between genotypes when fed a normal or high sucrose diet, suggesting that *dArc^-/-^* flies have similar fat mass and do not have a higher performance index because of lower energy stores (Extended Data Fig. 4i).

Sated flies are less motivated to show a preference for an odor they have learned to associate with a sucrose reward, as the expression of such appetitive memories is controlled by hunger state^18^. It is conceivable that the neuronal mechanisms that repress appetitive memories when flies are sated are defective in *dArc^-/-^* flies, leading to their enhanced performance in the sucrose reward conditioning paradigm. To explore this possibility, we trained *dArc^+/+^* and *dArc^-/-^* flies using sucrose rewards and tested their preference for the conditioned odor when flies were sated (Extended Data Fig. 6a). As expected, *dArc^+/+^* flies expressed appetitive memories only when starved. Similarly, *dArc^-/-^* flies expressed appetitive memories only when starved, suggesting that the mechanisms controlling the repression of appetitive memories in response to hunger state are intact.

### *dArc^-/-^* flies overvalue different types of sugar reward

Appetitive associative memories are formed when the positive value of a stimulus, such as a sugar reward, is associated with a neutral stimulus, such as an odor, through learning. The higher performance indices observed with *dArc^-/-^* flies may result from defects in the valuation of the sugar reward. Flies value sugar rewards based on both taste and nutritional content: sugars that taste sweet but lack nutritional content are not valued as highly as those that taste sweet and provide calories. When used in a conditioning paradigm, sweet-tasting sugars without caloric value typically lead to weaker memories than sweet-tasting sugars that provide nutritional value^19^. We used two different experimental assays to test whether *dArc^-/-^* flies value sugar rewards differently than *dArc^+/+^* flies. In the first set-up, we trained flies using arabinose, a sugar that tastes sweet but has no caloric value, and tested their preference for the conditioned odor either immediately after training or 30 minutes later. *dArc^-/-^*flies showed enhanced learning compared to *dArc^+/+^* flies when trained with arabinose, forming memories that were almost as strong as those formed with sucrose (Fig. 3a and Extended Data Fig. 6b). In the second assay, we trained flies using lower concentrations of sucrose — 10 mM, 100 mM and 250 mM. Only *dArc^-/-^*flies were able to learn at all concentrations and significantly outperformed *dArc^+/+^* flies when trained with 250 mM sucrose (Fig. 3b and Extended Data Fig. 6c). These experiments show that *dArc^-/-^* flies outperform *dArc^+/+^*flies when trained with sugar rewards that have no or low caloric value, suggesting an overvaluation of sugar reward.

**Figure 3.**
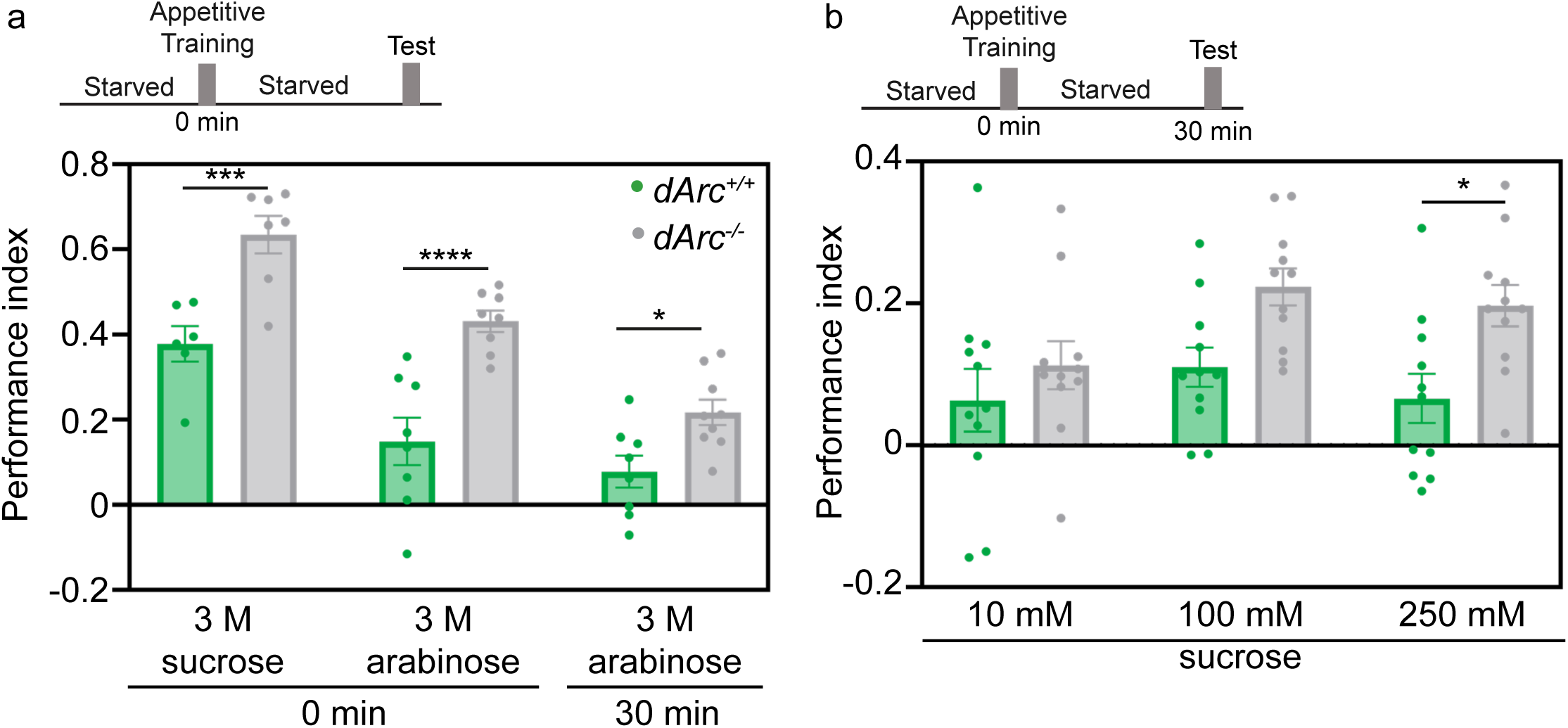
dArc^-/-^ flies overvalue sugar rewards. (a) Flies were starved for 20 hours and were subjected to an appetitive association paradigm using sucrose or the non-caloric sugar arabinose. The flies were then tested after 2 minutes for their immediate learning or 30 minutes for their short-term memory. (n ≥ 6 groups of flies, Two-way ANOVA). (b) Flies were starved for 20 hours and were subjected to an appetitive association paradigm using different concentrations of sucrose and tested after 30 minutes. (n ≥ 7 groups of flies, Two-way ANOVA). Subjects were mixed sex. Bar charts are presented as means ± s.e.m.

### γ5 dopaminergic neurons in *dArc^-/-^* flies show aberrantly strong responses to sucrose

In *D. melanogaster*, *a*ssociative learning takes place in the mushroom body^20,21^. The Kenyon cells that make up the mushroom body encode odor identity and are connected to different mushroom body output neurons that mediate either attraction or aversion to specific stimuli. Reward valence is encoded by dopaminergic neurons, where positive stimuli activate the protocerebral anterior medial (PAM) neurons^22,23^. Sugar rewards activate the PAM neurons that innervate the β’2 and γ5 compartments of the mushroom body, while water rewards activate those innervating the γ4 compartment^23^. The sugar valuation defects in *dArc^-/-^* flies may be due to hyperactivation of the β’2 and γ5 PAM neurons in response to sugar. To test this possibility, we recorded sugar-evoked activity in starved flies using calcium imaging and two-photon microscopy. We used the *R58E02-GAL4* transgene to target expression to the PAM neurons and the *UAS-GCaMP* transgene to record odor-evoked calcium transients^24,25^. Calcium transients were recorded in the β’2, γ4 and γ5 compartments following stimulation with 100 mM sucrose solution. We observed weak stimulus-evoked responses in the γ4 compartment, which were similar in magnitude in *dArc^+/+^* or *dArc^-/-^* flies (Fig. 4a). In the β’2 compartment, we observed strong stimulus-evoked responses, but there was no significant difference between *dArc^+/+^* and *dArc^-/-^*flies (Fig. 4b). In contrast, we observed strong stimulus-evoked responses in the γ5 compartment, with significantly higher responses in *dArc^-/-^* flies compared to *dArc^+/+^* flies (Fig. 4c). These experiments show that *dArc^-/-^* flies have functional defects in a restricted subset of dopaminergic neurons — the γ5 PAM neurons — that encode the value of sugar rewards.

**Figure 4.**
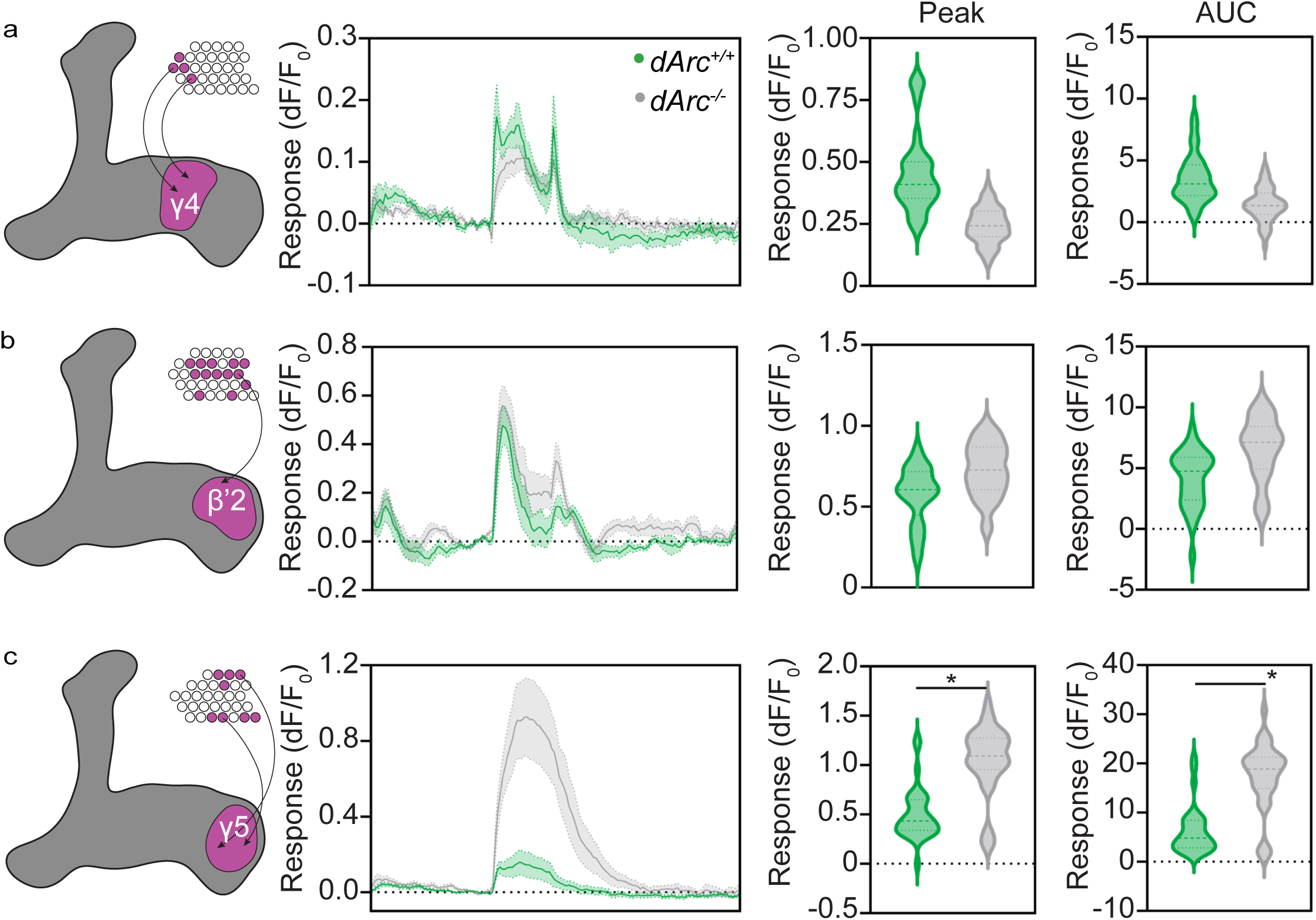
dArc^-/-^ flies have enhanced sugar responses in γ5 dopaminergic PAM neurons. (a) Starved flies expressing *UAS-GCaMP7f* in the protocerebral anterior medial dopaminergic neurons (*R58E02-GAL4*) were subjected to sucrose and their responses were measured in the γ4 dendritic arbors of the mushroom body. Total (AUC) and peak response of each response is quantified. (n ≥ 7 flies, 12-58 trials, nested t*-*test). (b) Starved flies expressing *UAS-GCaMP7f* in the protocerebral anterior medial dopaminergic neurons (*R58E02-GAL4*) were subjected to sucrose and their responses were measured in the β’2 dendritic arbors of the mushroom body. Total (AUC) and peak response of each response is quantified. (n ≥ 5 flies, 8-34 trials, nested t*-*test). (c) Starved flies expressing *UAS-GCaMP7f* in the protocerebral anterior medial dopaminergic neurons (*R58E02-GAL4*) were subjected to sugar and their responses were measured in the γ5 dendritic arbors of the mushroom body. Total (AUC) and peak response of each response is quantified. (n ≥ 10 flies, 16-36 trials, nested t*-*test). Subjects were mated females. Line graphs are presented as means ± s.e.m. Violin plots show the median (dashed line) and quartiles (dotted line).

### Capsid formation is necessary for dArc1 function in sucrose reward learning

dArc1 is expressed in a small number of neurons, most of which are serotonergic, but is not expressed in dopaminergic PAM neurons (Fig. 1 and Extended Data Fig. 2c,d). However, the defects we observed concern specific dopaminergic neurons, particularly the γ5 PAM neurons, raising the question of how dArc1 might exert its effect on these neurons. Previous studies have shown that dArc1 can self-assemble into virus-like capsids that are released in extracellular vesicles^4,3,6^. These observations suggest that dArc1 may affect dopaminergic neurons non-cell autonomously through the release of capsids by serotonergic neurons. To explore this possibility, we designed a dArc1 variant that cannot assemble into capsids (dArc1^ι1capsid^) (Fig. 5a) and tested whether rescuing the expression of *dArc1* or *dArc1^ι1capsid^* in serotonergic neurons could restore the enhanced learning phenotype observed in *dArc^-/-^*flies to *dArc^+/+^* levels. Purified dArc1^ι1capsid^ protein does not form capsids or oligomers (Fig. 5b-c, Extended Data Fig. 7a-c).

**Figure 5.**
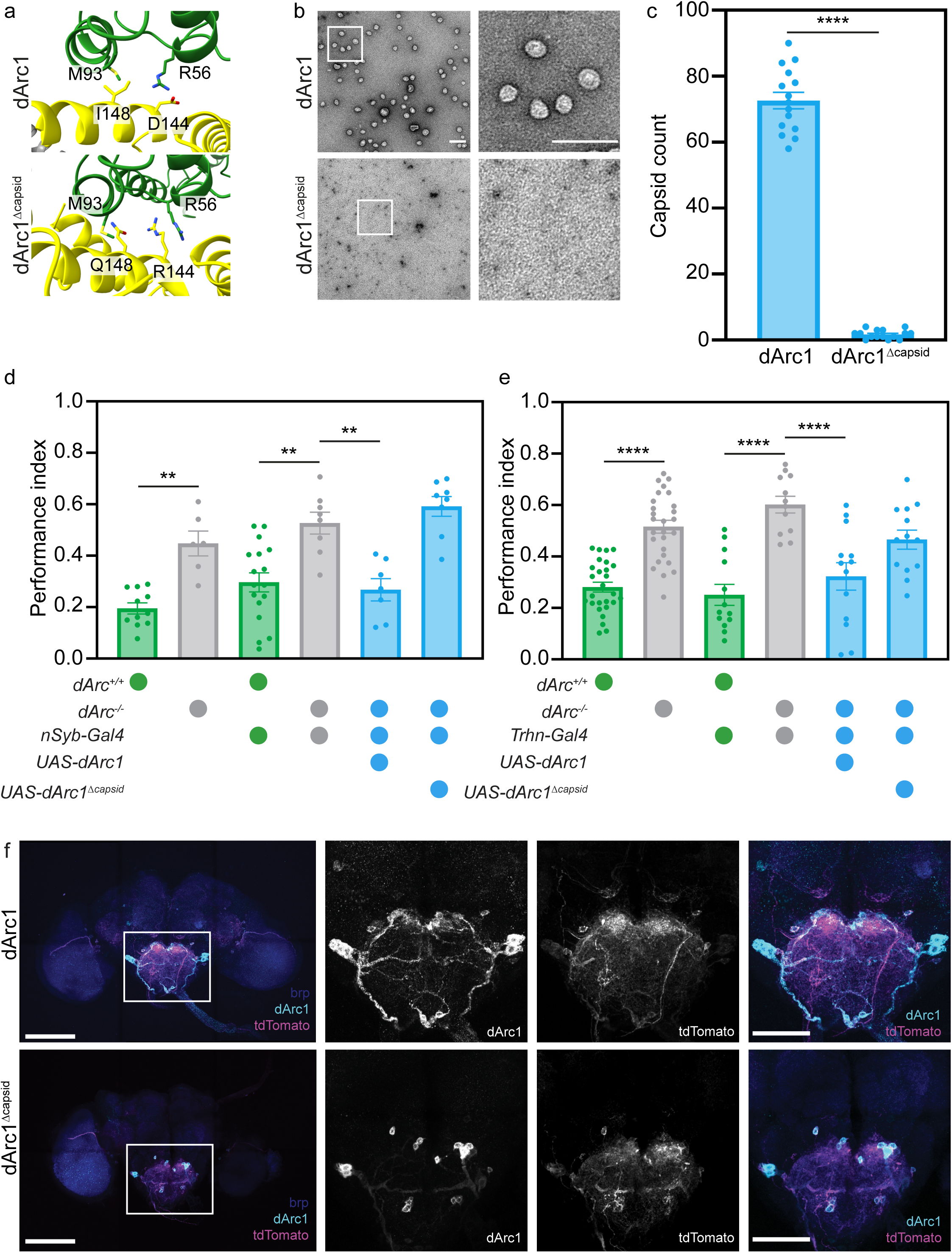
dArc1 capsids regulate sugar reward learning. (a) Visualization of the capsomer interface of two individual dArc1 proteins (green and yellow) indicating the introduced mutations. Colors on the displayed amino acid side chains indicate oxygen (red), nitrogen (blue), and sulfur (yellow). (b) Negative stain EM of dArc1 or dArc1^Δcapsid^ proteins. (c) Mean capsid counts from an electron microscope grid (n = 15 grid locations, Unpaired t*-*test) (d) Flies starved for 20 hours expressing either dArc1 or dArc1^Δcapsid^ in all neurons (*nSyb-GAL4*) were subjected to an appetitive association paradigm using sucrose and tested after 30 minutes. (n ≥ 6 groups of flies, One-way ANOVA). (e) Flies starved for 20 hours expressing either dArc1 or dArc1^Δcapsid^ in all serotonergic neurons (*Trhn-GAL4*) were subjected to an appetitive association paradigm using sucrose and tested after 30 minutes. (n ≥ 12 groups of flies, One-way ANOVA). (f) The brains of *dArc1/2^-/-^*flies expressing dArc1 and dArc1^Δcapsid^ proteins in all serotonergic neurons (*Trhn-GAL4*) labeled with *UAS-tdTomato* were immunostained using antibodies against the neuropil marker brp (blue), dArc1 (cyan), and mtdTomato (magenta). Shown is the anterior side of the brain and a zoom in of the suboesophageal zone. Scale bars are 100 µm. Subjects were mixed sex. Bar charts are presented as means ± s.e.m.

We tested whether rescuing the expression of *dArc1* or *dArc1^ι1capsid^* could restore the enhanced learning phenotype observed in *dArc^-/-^* flies to *dArc^+/+^* levels. We generated *UAS* transgenes encoding either *dArc1* or *dArc1^ι1capsid^*. We targeted expression of either transgene to all neurons using the *nSynaptobrevin-GAL4* transgene (*nSyb-GAL4*) or we targeted expression specifically to serotonergic neurons using the *tryptophan-hydroxylase-GAL4* (*Trhn-GAL4*). Rescuing *dArc1* in all neurons restored the learning phenotype observed in *dArc^-/-^* flies to *dArc^+/+^*levels (Fig. 5d, Extended Data Fig. 8a). In contrast, expression of *dArc1^ι1capsid^* did not affect the enhanced learning performance of *dArc^-/-^* flies, suggesting that capsid formation is necessary for *dArc1* function in appetitive learning. Similarly, we found that expression of *dArc1*, but not *dArc1^ι1capsid^*, in serotonergic neurons was sufficient to restore the learning phenotype observed in *dArc^-/-^* flies to *dArc^+/+^* levels (Fig. 5e). Expression of these transgenes in *dArc^+/+^* flies had no effect on learning performance (Extended Data Fig. 8b).

We confirmed the expression of both the *UAS-dArc1* and *UAS-dArc1^ι1capsid^* transgenes in neurons using the dArc1 antibody, which also recognizes dArc1^ι1capsid^ (Fig 5f and Extended Data Fig. 8a). Notably, the expression patterns differed between *dArc^-/-^*;*Trhn-GAL4*;*UAS-dArc1* and *dArc^-/-^*;*Trhn-GAL4*;*UAS-dArc1^ι1capsid^* flies. In *dArc^-/-^*;*Trhn-GAL4*;*UAS-dArc1* flies, dArc1 staining was distributed throughout serotonergic neurons, with distinct puncta observed along neuronal processes. In *dArc^-/-^*;*Trhn-GAL4*;*UAS-dArc1^ι1capsid^* flies, dArc1 staining was confined to the somata, suggesting that trafficking or secretion of dArc1^ι1capsid^ is disrupted.

## DISCUSSION

Our genetic and neurobiological experiments show that the retrotransposon-derived gene *dArc1* regulates associative learning by dampening the valuation of sugar rewards. Flies lacking *dArc1* overvalue sugar, leading to enhanced learning in an appetitive conditioning paradigm. This behavior is accompanied by hyperactivation of the γ5 PAM neurons in response to sucrose, suggesting that *dArc1* may regulate reward valuation by specifically modulating the activity of these neurons. Expressing *dArc1* in all serotonergic neurons corrected the enhanced learning phenotype in *dArc^-/-^*flies, while expression of a capsid assembly-deficient variant did not. These findings highlight the importance of dArc1 capsids, suggesting that this property is essential for its role in sugar valuation and associative learning, likely through intercellular signaling.

Such an enhanced learning phenotype is generally not known from studies of *Drosophila* neurogenetics and this may seem counterintuitive — why would the fly brain require a mechanism to suppress learning? However, this may make sense in the light of the nutritional physiology of *D. melanogaster*: Adult flies feed on fermenting fruit — an ephemeral food source that is rich in sugars — but also require proteins, provided by yeast colonizing fruit, in particular during egg production^26^. Excess carbohydrate consumption in *D. melanogaster* was shown to disrupt metabolic homeostasis in adults, impairing reproductive success and reducing lifespan^27–29^. By dampening dopaminergic reward signaling in response to sugar, Arc1 signaling may ensure that flies do not become overly dependent on sugar-rich food sources and continue to seek out a balanced diet. By contrast, when sugar is scarce, and Arc signaling down-regulated, carbohydrate rich food sources can be learned easily.

*dArc* and *mammalian Arc* (*mArc*) have an unusual evolutionary origin: they are independently derived from Ty3-type retrotransposons and encode proteins that form virus-like capsids that can travel between cells^3,4^. Here, we show that capsid formation is necessary for *dArc1* function in the fly brain. This raises the question of how this unusual mechanism of neuromodulation regulates sugar reward valuation. One possibility is that dArc1 signaling may induce state-dependent changes in target neurons that alter the ability to induce mechanisms of synaptic plasticity known from studies of learning in flies^30^. As dArc1 is expressed in serotonergic neurons — including those in the suboesophageal zone, which play key roles in modulating internal states^31^ — capsid signaling may integrate nutritional information over time, promoting adaptive plasticity in the neuronal circuits involved in associative learning. Further elucidation of dArc1 signaling mechanisms and — in particular the nature and function of the RNA cargoes transferred — will help address these questions.

The *Arc* genes represent a striking case study of evolutionary innovation in genomes by repurposing mobile genetic elements to enable vital organismal functions through unusual virus-like molecular mechanisms. *Arc* genes are found in two evolutionarily very successful lineages of animals: in tetrapod vertebrates, which count around 30,000 species, and in schizophoran flies, which count over 55,000 species^32,33^. Although this phylogenetic distribution indicates two independent integration events, both *dArc1* and *Arc* function in neuronal plasticity and appear to employ a similar capsid-dependent mechanism of signaling^3,4^. As transposon-derived sequences make up large swaths of animal genomes — over 50% of the human genome — Arc genes provide a window into the function and evolution of this ‘dark matter of the genome’^34^.

## ACKNOWLEDGMENTS

We thank members of the Caron and Shepherd laboratories for comments on the manuscript and discussions of the project. We are grateful to Dr. Adrian Rothenfluh for meaningful discussions on throughout the development of this project. We extend our heartfelt thanks to Dr. Florian Maderspacher for entertaining countless discussions on Arc and offering invaluable editorial insights. We also want to acknowledge Dr. Simon Erlendsson (University of Copenhagen) and Dr. John Briggs (Max Planck Institute of Biochemistry) for their help designing *dArc1^ι1capsid^*; Dane Larsen for the initial characterization of the dArc1 antibody; Adam Lin and Sylvia Yang for preparation of the standard cornmeal agar medium; Hayley Smihula and Lia Beatty for assistance with general laboratory concerns; the Cell Imaging Core at the University of Utah for use of the Zeiss LSM 880 microscope; and the Electron Microscopy Core at the University of Utah for use of the ThermoFisher JEOL-JEM 1400 microscope.

This work was initially funded by a Seed Grant awarded by the Neuroscience Initiative at the University of Utah (S.J.C.C., M.M.M. and J.D.S. and has been funded thereafter by grants from the National Institute for Neurological Disorders and Stroke (R01 NS 106018, R01 NS 1079790 and R01 NS 115716) and the National Science Foundation (IOS 2042397). Further financial support was provided by the Research Scholar Award (M.S.J. and B.D.M.), the University Research Opportunities Program (M.S.J.), the Chan-Zuckerberg Initiative (Ben Barres Early Acceleration Award; J.D.S.), the Jon M. Huntsman Presidential Endowed Chair Fund (J.D.S.) and the Georges S. and Dolores Eccles Foundation (S.J.C.C.).

## AUTHOR CONTRIBUTIONS

S.B., J.D.S. and S.J.C.C. conceived the project. S.B., J.D.S. and S.J.C.C. wrote the manuscript with input from all other authors. S.B. generated the UAS transgenic lines, performed and analyzed behavioral experiments (conditioning paradigms, feeding assay), performed expression analyses by qPCR and western blot, and generated the *dArc^-/-^* flies. M.S.J performed and analyzed behavioral experiments (conditioning paradigms). A.V.D. collected and analyzed the calcium imaging data. B.D.M. performed expression analyses by immunocytochemistry. K.R.S. performed protein purification and electron microscopy. A.R.B. performed expression analyses by immunocytochemistry and behavioral experiments (negative geotaxis assays, feeding assays). H.S. performed behavioral experiments (proboscis extension reflex), electrophysiological recordings, and metabolic assays under the supervision of M.D. J.E. and M.M.M. generated the *dArc^+/+^* and *dArc^-/-^* flies. J.D.S and S.J.C.C obtained funding for the project.

## MATERIALS AND METHODS

### Fly stocks

Flies (*Drosophila melanogaster*) were fed on standard cornmeal agar medium and raised in a controlled environmental chamber (Percival Scientific, Inc.) that maintains a temperature of 25°C and 60% humidity under a 12 hours light-12 hours dark cycle. Crosses were set up and reared under the same conditions. The strains used in this study are described in the table below.

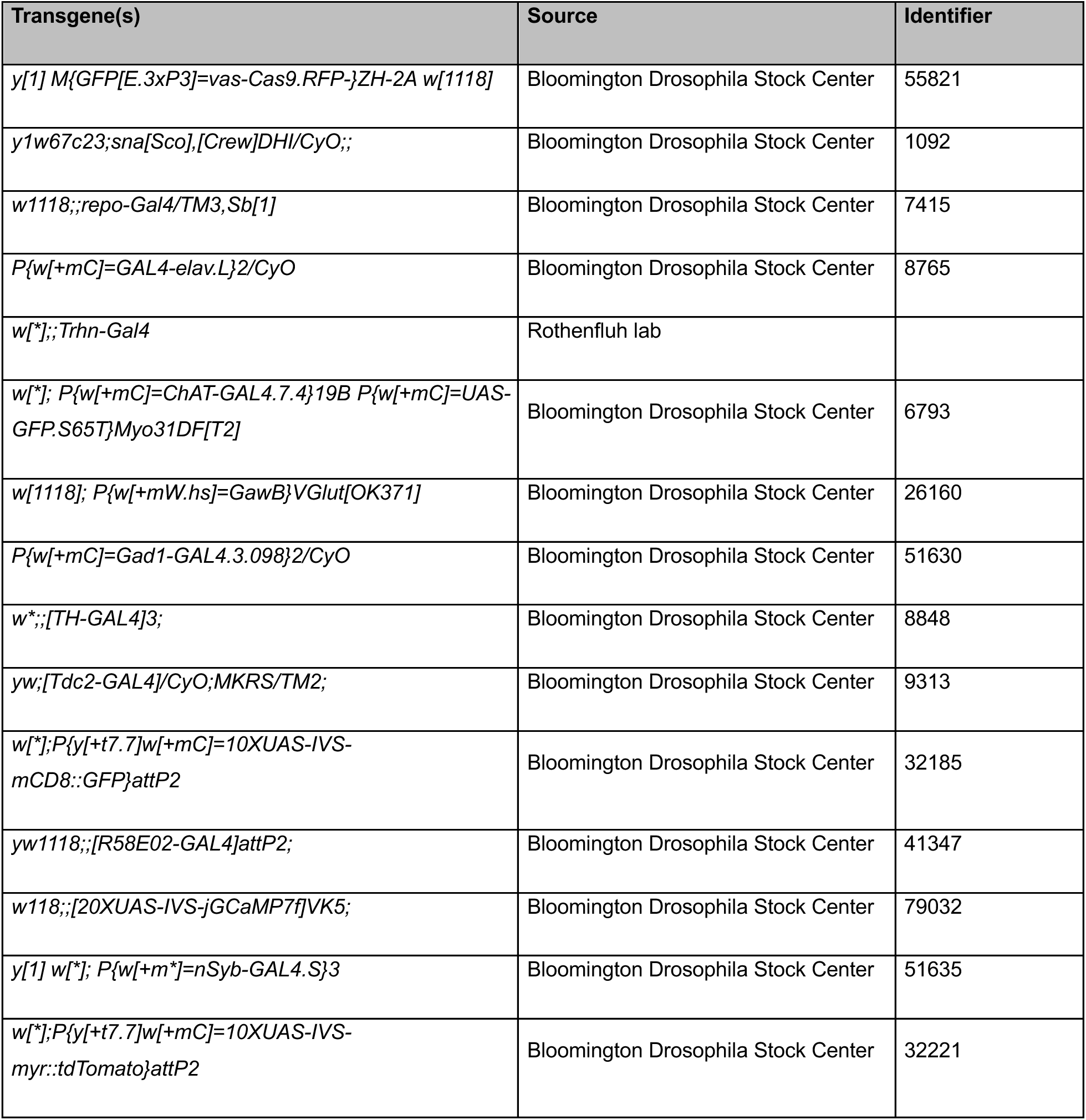

### Generation of *dArc^-/-^* flies

*dArc^-/-^* flies were generated using CRISPR/Cas9-mediated genomic engineering, following an established protocol^35,36^. Briefly, two guide RNAs (gRNAs) were designed to target various regions of *dArc1* and *dArc2* (gRNA1-2) and were evaluated for specificity and off-target effects using the CRISPR Target Finder tool (http://targetfinder.flycrispr.neuro.brown.edu/). These gRNA sequences were cloned into the pU6-BbsI-chiRNA plasmid and sequence-verified. Homology arms of about 1 kb flanking the Cas9 cut sites were synthesized as gene blocks (IDT) and cloned into the pHD-DsRedattP plasmid, followed by sequence verification. The gRNAs and homology repair plasmids were co-injected into *vas-Cas9* transgenic embryos (Bestgene). Adult flies were backcrossed to *w^11^*^18^ flies. Mutant chromosomes were identified using the dsRed marker — which was under the *3xP3* promoter and therefore visible in adult eyes — and isolated using the balancer chromosome *CyO*. The resulting *dArc^-/-;DsRed^*flies were crossed to flies expressing Cre recombinase (*Cre*) to excise the DsRed marker, yielding the *dArc^-/-^* flies used throughout this study. *dArc^-/-^* flies were validated by PCR using primers flanking the targeted region. The control line, *dArc^+/+^* flies, was established by isolating the second chromosome from *vas-Cas9* flies.

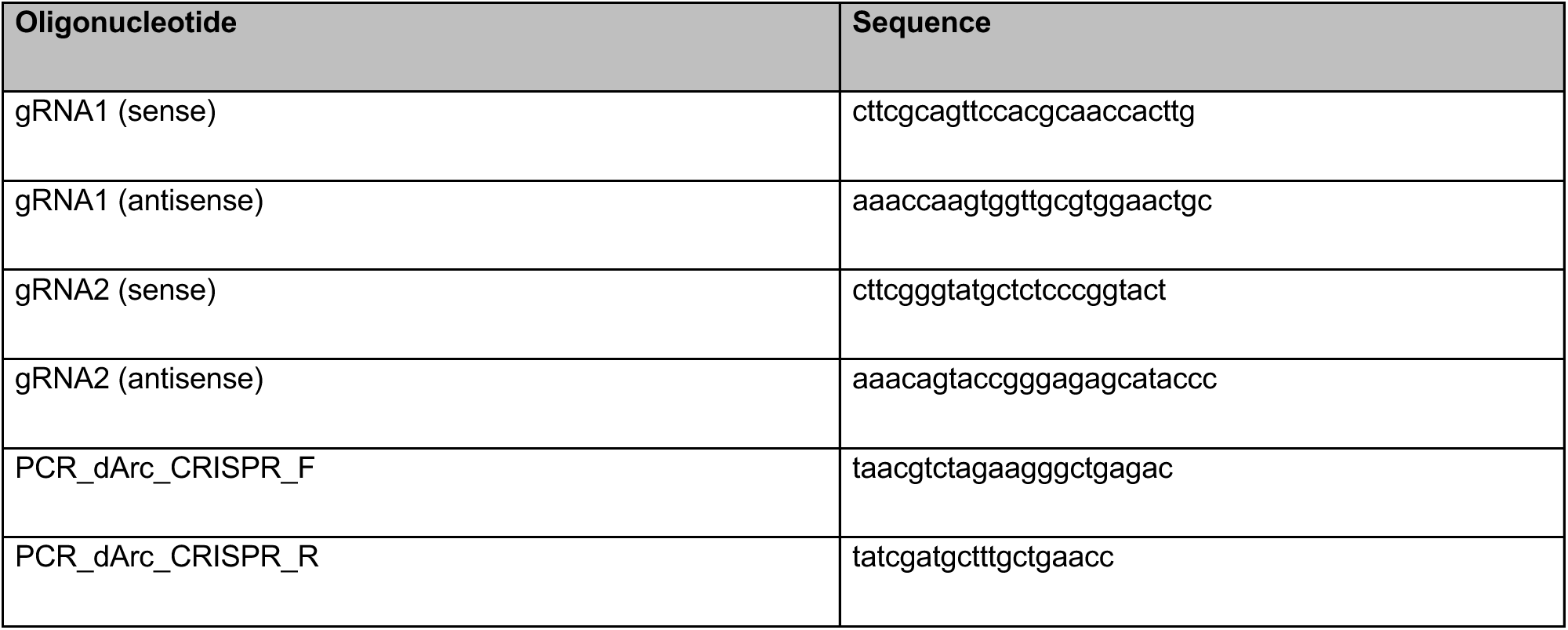

### Generation of transgenic flies

The *UAS-dArc1* transgene was generated using Gibson cloning, following previously described protocols^37,38^. Briefly, *dArc1*, including its 3’-UTR, was cloned into the pUASt-attB plasmid (DGRC Stock 1419; https://dgrc.bio.indiana.edu/stock/1419; RRID:DGRC_1419) using primers with 30 base pairs homology arms. The construct was sequence-verified. The plasmid was inserted into the attPVK00033 site using PhiC31-mediated recombination^39^. Transgenic flies were identified by the presence of the mini-white marker and isolated using the balancer chromosome *TM6B*. Point mutations were introduced in the pUASt-attB-dArc1 plasmid to generate the *UAS-dArc1^Dcapsid^* transgene (Genescript). The mutated plasmids were sequence-verified. The plasmid was inserted in the attPVK00033 site using PhiC31-mediated recombination. Transgenic flies were identified by the presence of the mini-white marker and isolated using the balancer chromosome *TM6B*.

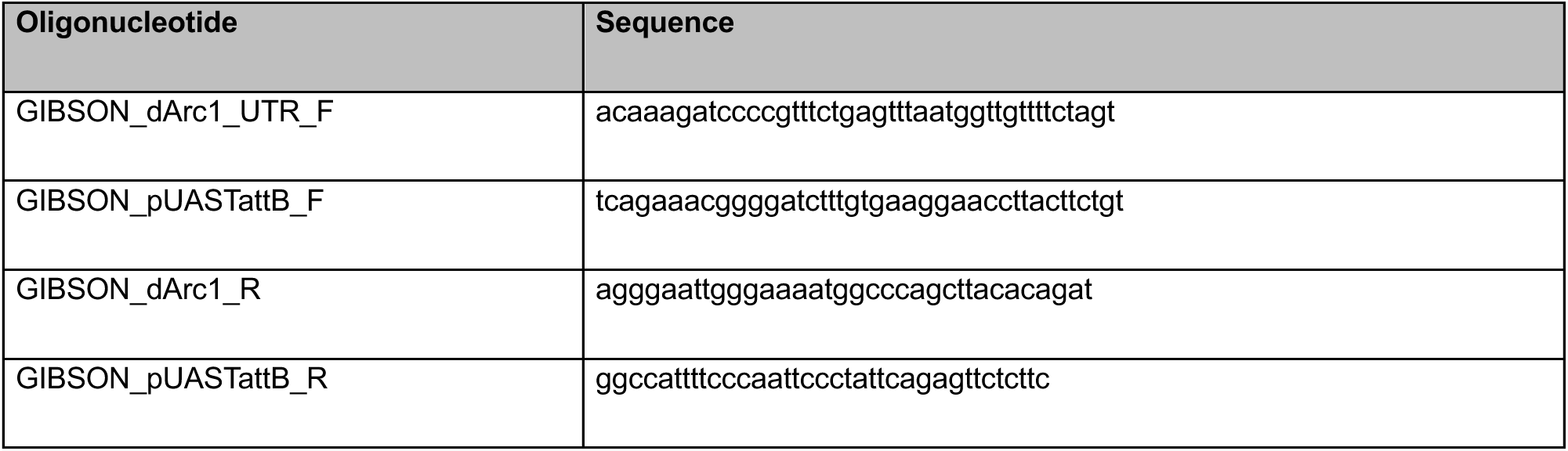

### Generation of Arc1 and Arc2 antibodies

Custom dArc1 antibodies were generated by the Proteintech Group. *dArc1* was cloned into the pGEX-4T1 plasmid. The GST-dArc1 fusion protein was purified and used to immunize two rabbits. Four booster immunizations were administered before the final bleed was collected from each rabbit. The custom antibodies were then affinity-purified using 6xHis-dArc1 and stored in 10% glycerol. Custom dArc2 antibodies were generated by ThermoFisher. A dArc2 peptide was synthesized (CKTFRELLDRGRTVERTRH) and used to immunize two guinea pigs. Three booster immunizations were administered before the final bleed was collected from each guinea pig. The custom antibodies were then affinity-purified using the dArc2 peptide and stored in 1x PBS with 0.05% NaN3.

### Immunohistochemistry

The brains of one to five day-old flies were dissected in saline and fixed in PFA for 45 min at room temperature, washed five times in PBST at room temperature, blocked in PBST-NGS for 30 min at room temperature, and incubated with different primary antibodies at 4°C overnight. Depending on the experiment, different antibodies were used at different concentrations (see table below). On the following day, brains were washed four times in PBST and incubated in secondary antibodies at 4°C overnight. Different secondary antibodies were used (see table below). On the following day, brains were washed four times in PBST and mounted on a slide (Fisher Scientific) using the mounting media VECTASHIELD (Vector Laboratories Inc.). The neuropils were identified by comparing the confocal images with the adult brain template available in the FlyWire Codex (https://codex.flywire.ai/app/neuropils)^40,41^.

| Antibody | Concentration | Source |
| --- | --- | --- |
| Rabbit anti-dArc1 | 1:1000 | This study |
| AlexaFluor-546 goat-anti-rabbit | 1:500 | Life technologies |
| AlexaFluor-488 goat-anti-rabbit | 1:500 | Life technologies |
| Mouse anti-brp (nc82) | 1:20 | DSHB |
| AlexaFluor-633 goat-anti-mouse | 1:500 | Life technologies |
| AlexaFluor-546 goat-anti-mouse | 1:500 | Life technologies |
| Rat anti-tdTomato (16D7) | 1:500 | Kerafast |
| Alexa Fluor 568 goat-anti-rat | 1:500 | Life technologies |

### Confocal image acquisition and analysis

All confocal images were collected on LSM880 confocal microscope (Zeiss). For imaging whole brains, each sample was imaged using a Plan-Apochromat 40X/1.3 Oil M27 objective lens. First, the entire brain was divided into six tiles, each tile was imaged separately (voxel size = 0.21 μm by 0.21 μm by 0.44 μm) and then stitched together using the Stitch function of the Zen microscope software (Zeiss). All confocal images were handled using the Fiji software^42^. All figure panels are maximum intensity projection of confocal stacks or sub-stacks.

### Conditioning paradigms

Aversive and appetitive conditioning paradigms were performed following previously described protocols that use a T-maze^15,16,43,44^. Both paradigms consist of a training phase and a testing phase. Each experiment was carried out on a group of 60 to 100 flies. In general, flies were selected prior to an experiment without sedation, except when selecting for the correct genotype. In such cases, flies were allowed to recover from anaesthesia for at least one full day. For the aversive paradigms, flies were kept on standard cornmeal fly food before the training phase. For the appetitive paradigms, flies were kept on 1% agar for 20 hours prior to the training phase. The T-maze was used to train and test flies. In this T-maze, odors (4-methylcyclohexanol or 3-octanol; Sigma-Aldrich) were delivered through an air-flow that was held stable at 0.750L per minute. Odors were diluted in light mineral oil (Sigma-Aldrich), and the approximate concentration used in each experiment was ∼1:10^-3^ concentration. Odor concentrations were optimized to achieve a balanced distribution of naïve flies between the two arms of the T-maze prior to the training phase.

In the short-term appetitive conditioning paradigm, starved flies were initially placed in a tube containing an air-dried piece of paper (US^-^) while exposed to an odor (CS^-^) for two minutes, followed by a stream of unscented air for 30 seconds. Flies were transferred to a different tube containing a piece of paper with air-dried sugar (sucrose or arabinose; US^+^) or a paper with dissolved sucrose in 1% agar (for different concentrations of sucrose; US^+^) while exposed to another odor (CS^+^) for two minutes, followed by a stream of unscented air for 30 seconds. Trained flies were returned to their original vial for the indicated time before the testing phase. In the long-term appetitive conditioning paradigm, flies were kept at 18°C before being tested.

In the short-term appetitive conditioning paradigm that used water as the reward, flies were desiccated by being kept in a tube with a piece of paper with air-dried sugar for 20 hours. Desiccated flies were then placed in a tube containing an air-dried piece of paper (US^-^) while exposed to an odor (CS^-^) for two minutes, followed by a stream of unscented air for 30 seconds. Flies were transferred to a different tube containing a wet piece of paper (US^+^) while exposed to another odor (CS^+^) for two minutes, followed by a stream of unscented air for 30 seconds. Flies were returned to their original vial for the indicated time before the testing phase.

In the short-term aversive conditioning paradigm, flies were placed in a copper-lined tube and exposed to an odor (CS^+^) while receiving twelve 1-second 90V shocks over one minute (US^+^), followed by a stream of unscented air for 45 seconds. Flies were then exposed to a second odor (CS^-^) without electric shocks (US^-^), followed by another stream of unscented air for 45 seconds. In the long-term aversive conditioning paradigm, flies underwent six training sessions, each consisting of twelve 1-second 90V shocks given over one minute, with a 15-minute interval between sessions. Flies were returned to their original vial for the indicated time before the testing phase.

During the testing phase for both the appetitive and aversive conditioning paradigms, trained flies were given the choice to enter one of two tubes in a T-maze: one scented with the conditioned odor previously paired with an appetitive reward or punitive shocks (CS^+^) and the other tube was scented with the conditioned odors that was not paired with an appetitive reward or punitive shocks (CS^-^); flies had two minutes to make a choice. The flies located in each tube were collected and placed at -80°C at the end of the experiment. The number of flies in each tube was counted, and the performance index was calculated using the following formula: 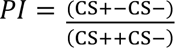. Flies that did not make a choice were discarded.

### Calcium imaging

Calcium imaging experiments were performed using similar methods as previously described^45^. Flies were immobilized on their backs using a piece of clear tape in an imaging chamber. Fine strands of tape were used to restrain the legs, secure the head, and immobilize the proboscis in an extended position. A dorsal imaging window was cut in the fly cuticle. Images were collected using a two-photon laser scanning microscope (Investigator, Bruker) equipped with an ultra-fast Ti:Sapphire laser (Ultra II, Coherent) modulated by pockel cells (Conoptics). The excitation wavelength was 925 nm. Emitted photons were collected with a GaAsP photodiode detector (Hamamatsu) through a 20X water-immersion objective (Olympus). Images were acquired using the microscope software (PrairieView, Bruker). Imaging planes were chosen for optimal dendritic signal of each dopaminergic compartment in the mushroom body. Images were acquired at 512 by 512 pixels with a scanning rate of ∼5 Hz.

Taste stimuli were delivered as drops to the labellum using a custom-built solenoid pinch valve system controlled by a MATLAB software through a data acquisition device (Measurement Computing). Solenoid pinch valves (Cole Parmer) were briefly opened (30-50 milliseconds) to create a small liquid drop at the end of a glass capillary. The capillary was positioned such that the drop made contact with the labellum. The drop was removed after five seconds by a vacuum line controlled by a solenoid pinch valve. Proper delivery of the drop was monitored using a side-mounted camera (FLIR Blackfly), which visualized the fly and the capillary. At least eight trials were conducted, with a minimum rest period of one minute between trials to avoid habituation. The taste stimulus used in all recordings was 100 mM sucrose. Only flies that displayed signs of health and activity — such as abdomen or proboscis movement — were included.

Calcium transients were analyzed using custom MATLAB (MathWorks) codes. Movement between individual images corrected, within and across trials for the same fly. Calcium transients (ΔF/ F0) were calculated by measuring the average fluorescence before and during stimulation (F0: four seconds prior to stimulus onset, F: three seconds during stimulus delivery). Data were pooled for each fly across trials and hemispheres for each imaged compartment.

### Design of the dArc1^ι1capsid^ variant

The structure of dArc1 capsids, previously resolved by cryo-electron microscopy, was used to design the dArc1^ι1capsid^ variant. To disrupt the tertiary interactions between pentamers and hexamers necessary for capsid assembly, without altering the structure of the monomeric subunits, the aspartic acid at position 144 was substituted with an arginine and the isoleucine at position 148 with a glutamine. dArc1 and dArc1^ι1capsid^ were expressed in *Escherichia coli* and purified using glutathione S-transferase (GST) affinity purification followed by size exclusion chromatography. The purified dArc1 protein eluted early in the void fraction, indicating the presence of high molecular weight species, such as capsids. In contrast, the dArc1^ι1capsid^ protein eluted in later fractions, suggesting a lack of high molecular weight species (Extended Data Fig. 7a-c). The void fractions was imaged using negative stain electron microscopy.

### Protein structure prediction

Protein structure predictions were performed using the Alphafold3 server (https://golgi.sandbox.google.com/about)^46^. The protein sequences of dArc1 and dArc1^Δcapsid^ were used. The molecule type was set at protein with five copies of the protein without any post-translational modifications. The output was visualized using the chimeraX software (https://www.rbvi.ucsf.edu/chimerax)^47^.

### Protein purification

GST-tagged proteins were purified from *Escherichia coli* Rosetta 2 BL21 competent cells as previously described^3^. Starter bacterial cultures were grown overnight at 37°C in LB medium supplemented with ampicillin and chloramphenicol. Large-scale 500 mL cultures were inoculated with the starter cultures in ZY auto-induction medium. These cultures were grown to an optic density of 600 nm of 0.6–0.8 at 37°C with shaking at 160 rpm, then shifted to 18°C with shaking at 180 rpm for 16–20 hours. Cultures were pelleted at 4000x*g* for 15 minutes at 4°C, and the resulting pellets were resuspended in 25 mL of lysis buffer (500 mM NaCl, 50 mM Tris, 5% glycerol, 1 mM DTT, pH 8.0) before being flash-frozen in liquid nitrogen. Frozen pellets were thawed in a 37°C water bath and brought to a final volume ratio of 1 g pellet:10 mL lysis buffer, supplemented with DNase, lysozyme, and a complete protease inhibitor cocktail (Roche).

Resuspended lysates were sonicated for six 45-second pulses at 90% duty cycle and then centrifuged at 21,000x*g* for 75 minutes. The cleared supernatants were incubated with pre-equilibrated GST Sepharose 4B affinity resin in a gravity flow column overnight at 4°C. GST-bound proteins were washed twice with 20 resin bed volumes of lysis buffer and then re-equilibrated in TBS (150 mM NaCl, 50 mM Tris, 1 mM EDTA, 1 mM DTT, pH 7.2) at room temperature. The proteins were cleaved from the GST resin by incubation with PreScission Protease (GE Healthcare) overnight at 4°C.

### Arc capsid assembly assay

Glutathione S-transferase-cleaved and purified dArc1 and dArc1^Dcapsid^ protein eluates were concentrated to 2.5 mL using Vivaspin 10 kDa MWCO ultrafiltration spin columns (Cytiva). Protein concentration was measured using the DC Protein Assay (BioRad). Equal concentrations of dArc1 or dArc1^Dcapsid^ proteins were then applied to a Superdex 200 pg HiLoad 16/600 size exclusion column in Tris-Buffered Saline. Peak fractions were identified by chromatogram analysis and Coomassie staining, pooled, and concentrated. Following size exclusion, negative stain electron microscopy grids were prepared for each sample at a protein concentration of 0.5 mg/mL.

### Negative stain electron microscopy

For all negative stain electron microscopy samples, copper 200-mesh grids (Electron Microscopy Sciences) were glow-discharged for 25 seconds in a vacuum chamber at 30 milliamperes. A 3.5-microliter sample was applied to the grid for 45 seconds. Grids were washed twice, for five seconds each, with 30-microliter water droplets, followed by one wash with one percent uranyl acetate on parafilm. Excess water and uranyl acetate were removed from the grid using filter paper, and a final droplet of uranyl acetate was added to the grids for 30 seconds. Excess uranyl acetate was then removed, and the grids were air-dried for 30 seconds. Imaging was performed using a JEOL-JEM 1400 electron microscope at 120 kilovolts. For each experiment, 15 images were taken of each sample, following a standardized pattern of grid squares. Arc capsids ranging from 30 to 50 nanometers were manually counted using ImageJ/FIJI.

### Statistical analysis

Measurements were taken from independent samples. All statistical analyses were performed using GraphPad Prism 10.3.1 (GraphPad Software, San Diego, CA). Individual tests for each dataset are reported within the figure legends. Reported bar charts show the mean and the error bars as s.e.m. Reported line charts show the mean and the error bars as s.e.m. Reported violin plots show the median (dashed line) and quartiles (dotted line). For all statistical analyses, significance was set at *p*-value of less than 0.05. In all figures the *p*-value is reported as \**p* < 0.05, \*\**p* < 0.01, \*\*\**p* < 0.001, \*\*\*\**p* < 0.0001. ns, non-significant.

---

**Extended data figure 1.**
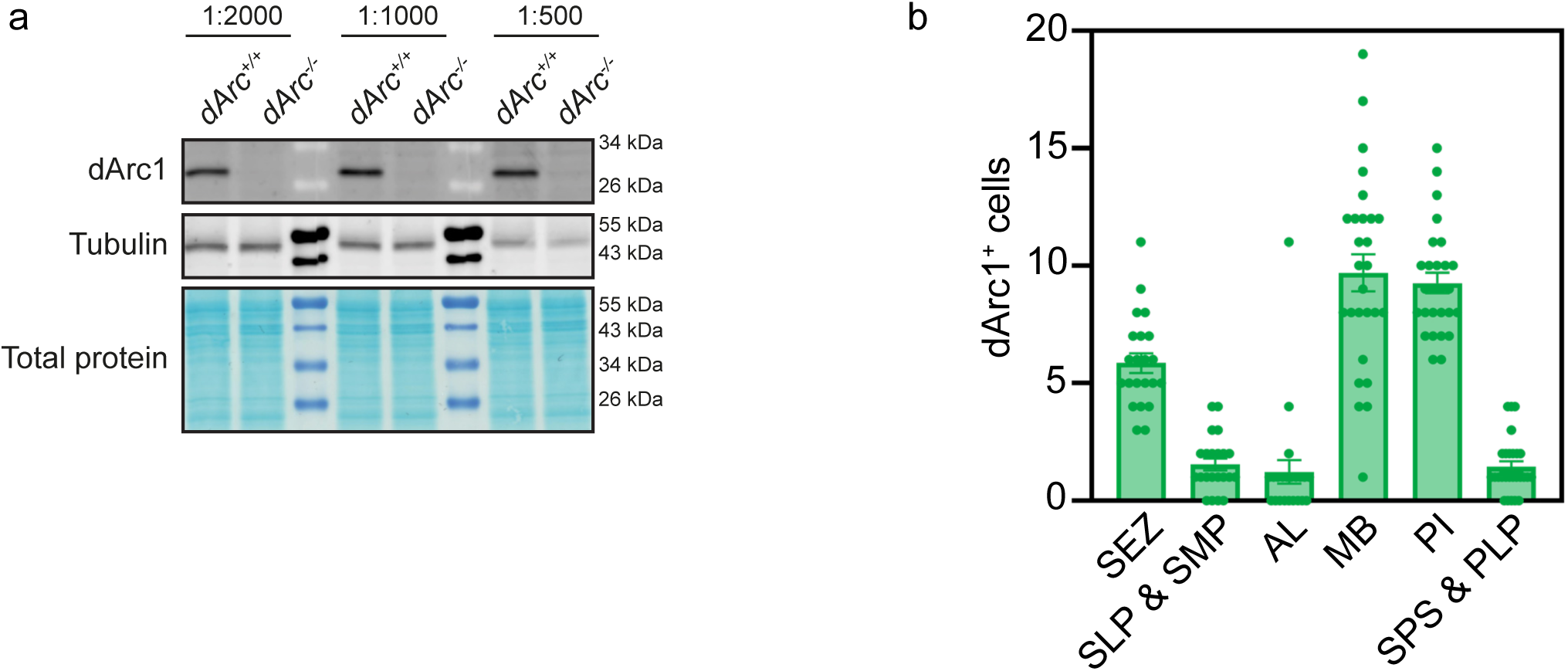
(a) The expression of dArc1 protein in the adult fly head was determined by western blot using a custom rabbit antibody. Tubulin and total protein were used as loading controls. (b) The localization of dArc1 positive cells was determined in the *dArc^+/+^*fly brain. SEZ, subeosophageal zone; SLP & SMP, superior lateral and superior medial protocerebrum; AL, antennal lobes; MB, mushroom body; PI, posterior inferior; SPS & PLP, superior posterior slope and posterior lateral protocerebrum.

**Extended data figure 2.**
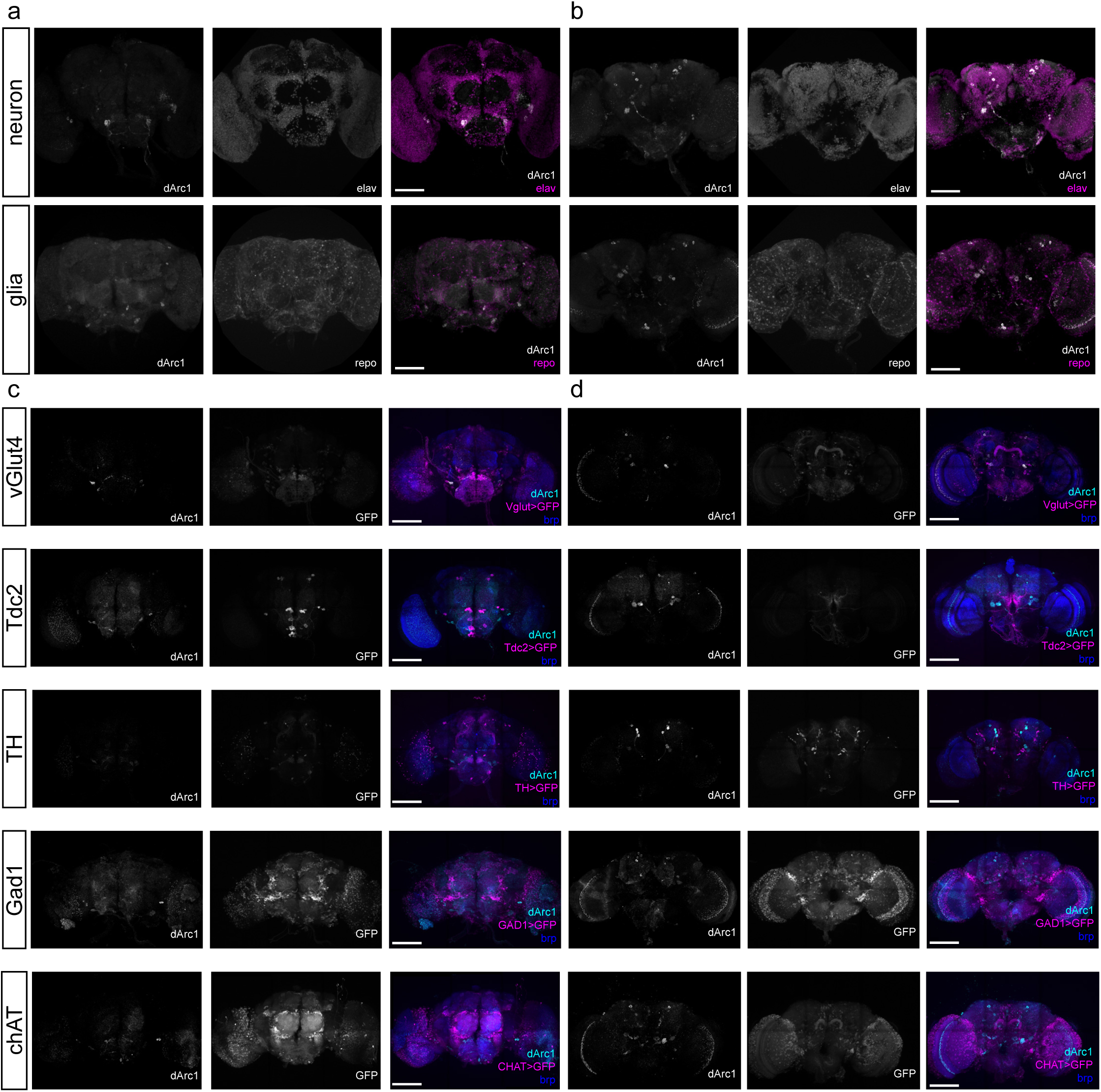
(a) The brains of *dArc^+/+^* flies were immunostained using antibodies against the neuronal transcription factor elav (magenta) or glial marker repo (magenta), and dArc1 (white). Shown are the anterior and (b) posterior views. Scale bars indicate 100 µm. (c) The brains of *dArc^+/+^* flies expressing UAS-GFP in specified neurotransmitter subtypes (magenta) were immunostained using antibodies against dArc1 (cyan) and a neuropil marker brp (blue). *vGlut4-GAL4*: glutamatergic, *Tdc2-GAL4*: octopaminergic, *TH-GAL4*: dopaminergic, *Gad1-GAL4*: GABAergic, *chAT-GAL4*: cholinergic neurons. Shown are the anterior and (d) posterior views. Scale bars indicate 100 µm.

**Extended data figure 3.**
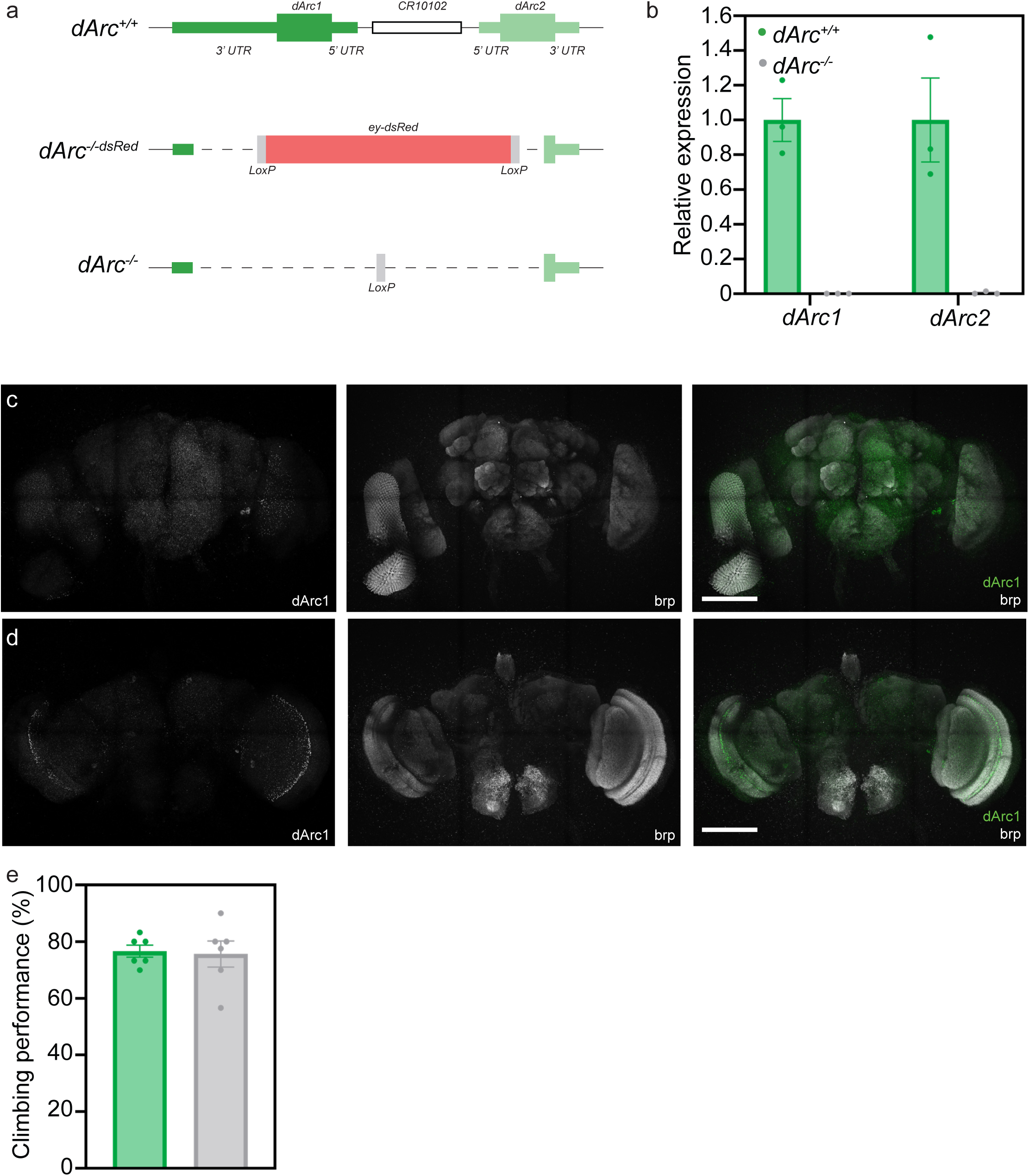
(a) Schematic overview of the dArc genetic locus showing *dArc1* on the minus strand and *dArc2* on the plus strand flanking the *CR10102* pseudogene. The three genes were replaced by an ey-DsRed cassette using independent gRNAs targeting *dArc1* and *dArc2*. The ey-DsRed cassette was then removed using Cre recombinase and the available LoxP sites. (b) The expression of *dArc1* and *dArc2* mRNA was determined by quantitative PCR (n = 3 replicates). (c) Anterior view of a *dArc^-/-^* fly brain that was immunostained with dArc1 and Brp antibodies. Scale bar represents 100 µm. (d) Posterior view of a *dArc^-/-^* fly brain that was immunostained with dArc1 and Brp antibodies. Scale bar represents 100 µm. (e) The percentage of flies crossing the 7cm mark during the negative geotaxis assay (n ≥ 6 groups of flies, Unpaired t-test). Subjects were female.

**Extended data figure 4.**
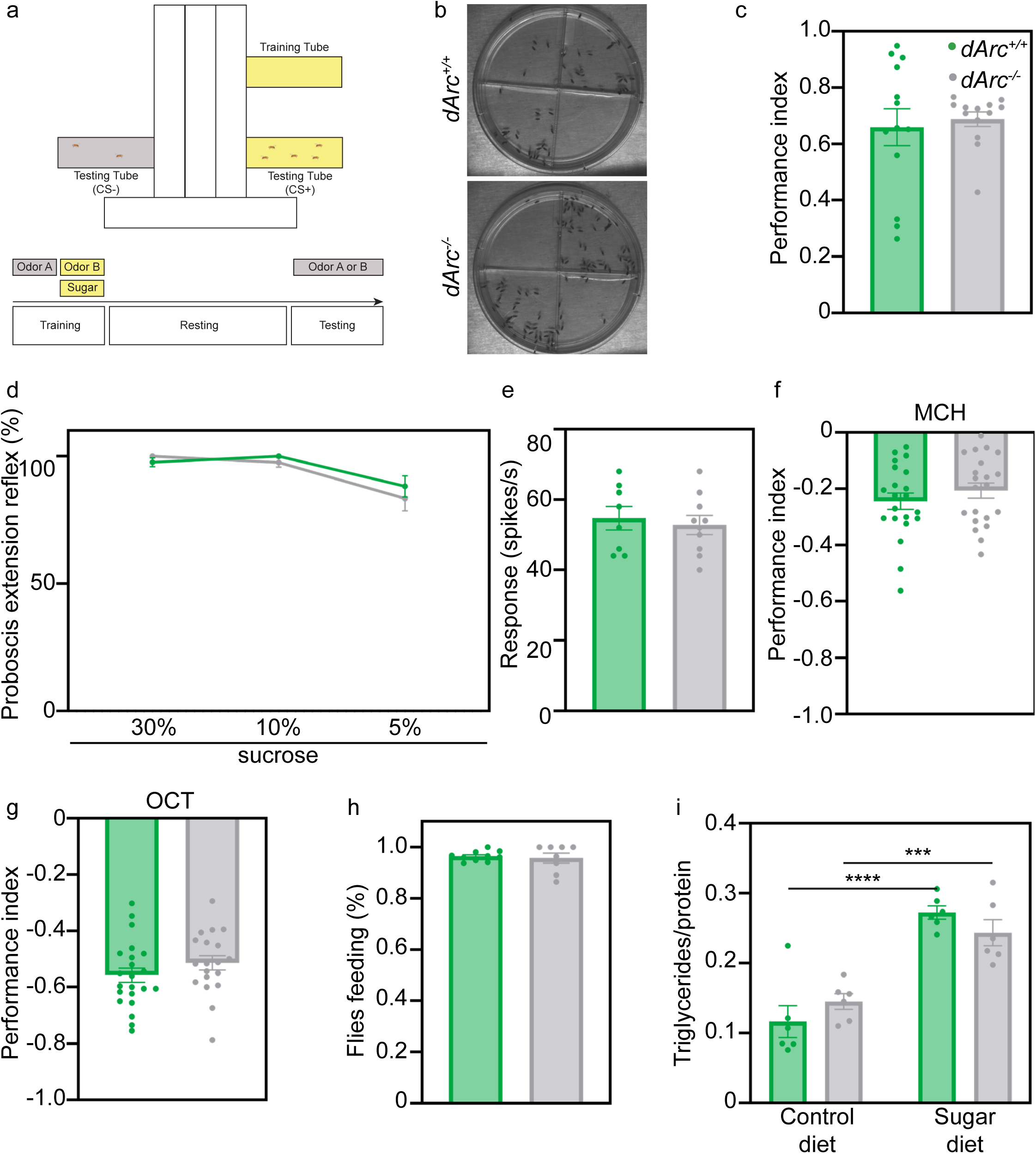
(a) A schematic of the *Drosophila* T-maze setup. Flies were initially exposed to an odor A without positive reinforcement (US-), followed by an odor B with a dried sucrose reward (US+). After a given rest period, flies were tested in a 2-choice assay between odor A (CS-) and B (CS+). (b) Acuity for sugar of starved *dArc^+/+^* and *dArc^-/-^* flies was tested using a place-preference assay. Representative images are shown. (c) The performance index for sugar acuity was determined for starved *dArc^+/+^* (green) and *dArc^-/-^* (grey) flies after 5 minutes (n ≥ 13 groups of flies, Unpaired t-test). (d) The percentage of *dArc^+/+^* (green) and *dArc^-/-^* (grey) flies showing a proboscis extension reflex was determined in response to different concentrations of sucrose (n ≥ 20 flies, Two-way ANOVA). (e) The electrophysiological responses of the sweet sensing sugar neurons of *dArc^+/+^* (green) and *dArc^-/-^* (grey) flies was determined after exposure to sucrose (n ≥ 8 flies, Unpaired t-test). (f) The odor acuity of *dArc^+/+^* (green) and *dArc^-/-^* (grey) flies for 4-methylcyclohexanol (MCH) (n = 21 groups of flies, Unpaired t-test) and (g) 3-octanol (OCT) was determined in a 2-choice assay for 2 minutes (n ≥ 19 groups of flies, Unpaired t-test). (h) The fraction of flies feeding on the sucrose reward was determined for 2 minutes (n ≥ 8 groups of flies, Unpaired t-test). (i) The metabolic responses of *dArc^+/+^* (green) and *dArc^-/-^* (grey) flies were determined after flies were fed a control or sugar diet. The ratio of triglycerides per protein is shown (n ≥ 6 groups of flies, Two-way ANOVA). Subjects were mixed sex. Bar charts are presented as means ± s.e.m.

**Extended data figure 5.**
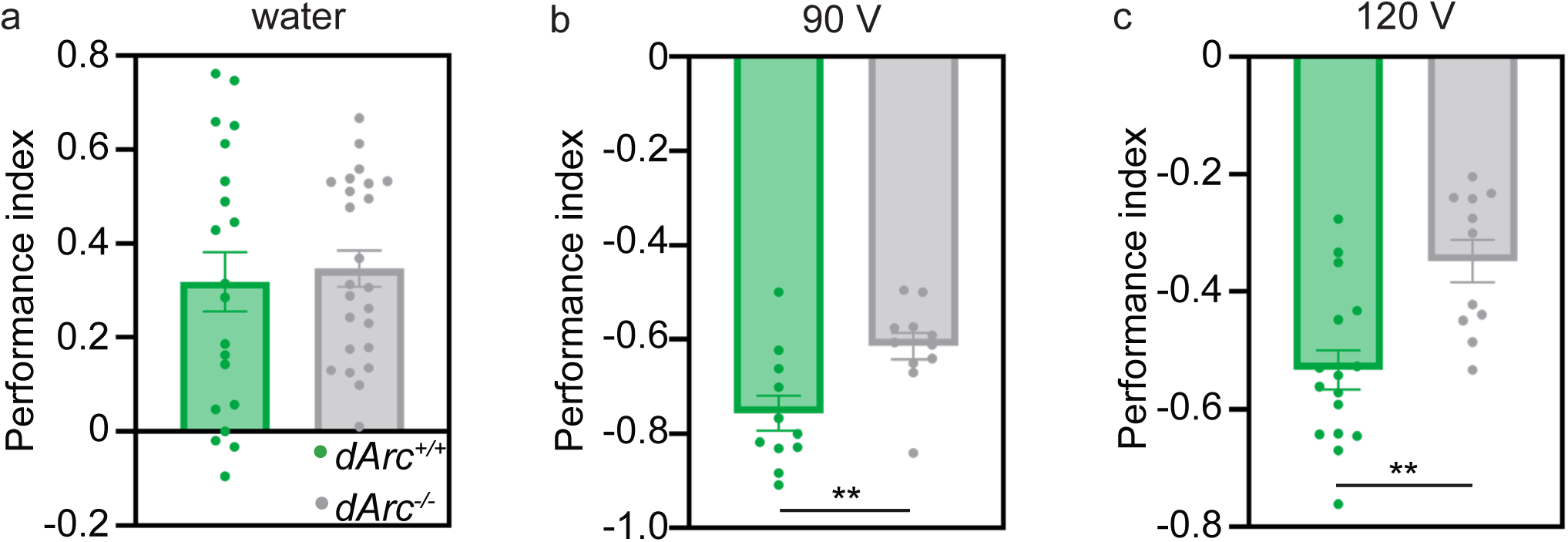
(a) The performance index for water acuity was determined for *dArc^+/+^* (green) and *dArc^-/-^* (grey) flies in a 2-choice assay (n ≥ 20 groups of flies, Unpaired t-test). (b) The performance index for electric shock acuity was determined for *dArc^+/+^* (green) and *dArc^-/-^* (grey) flies using 90V (n ≥ 11 groups of flies, Unpaired t-test) and (c) 120V shocks using a 2-choice assay (n ≥ 11 groups of flies, Unpaired t-test). Subjects were mixed sex. Bar charts are presented as means ± s.e.m.

**Extended data figure 6.**
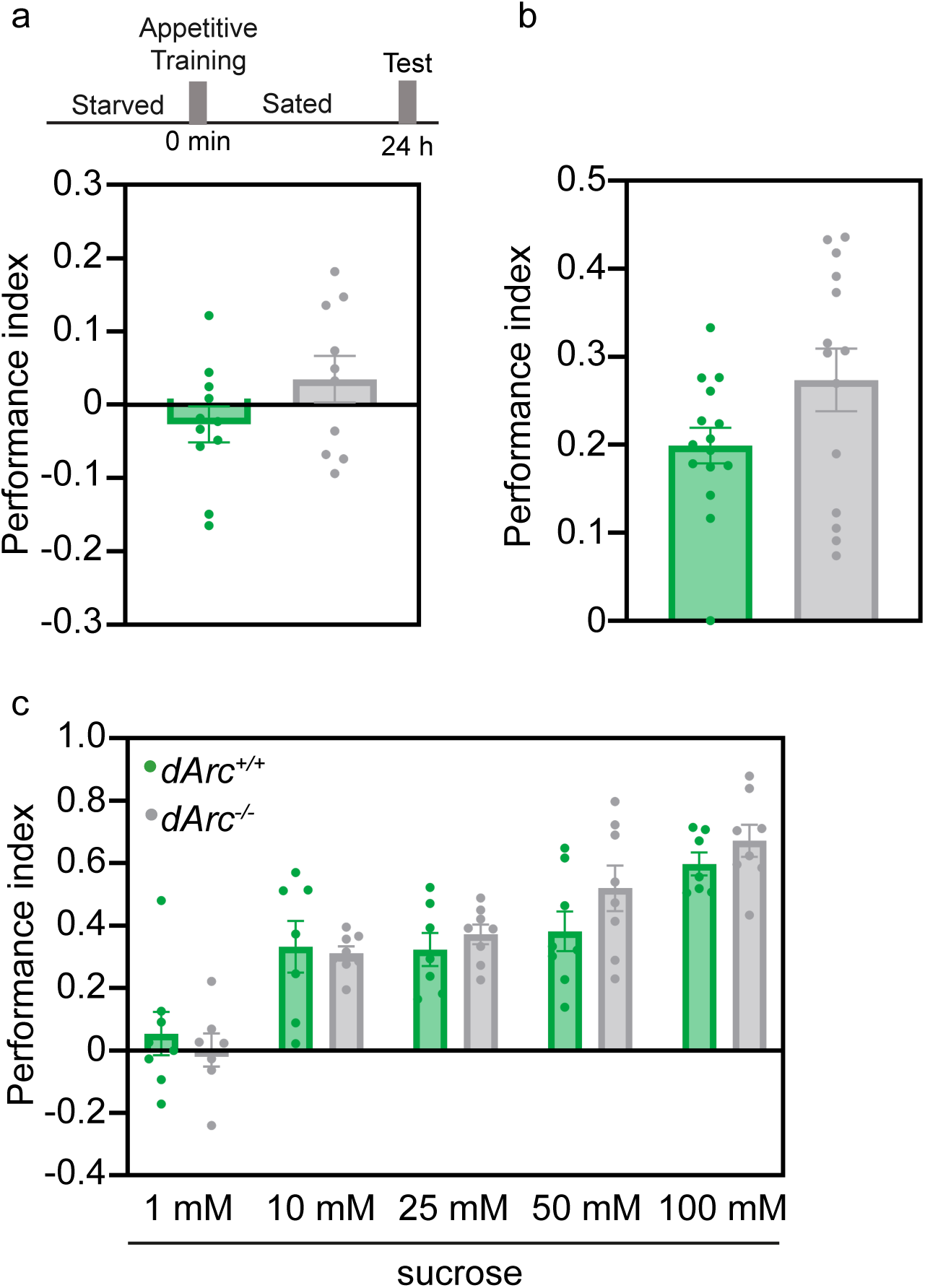
(a) Flies were starved for 20 hours and were subjected to an appetitive association paradigm using sugar. The flies were then fed ad libitum after training and tested after 24 hours. (n ≥ 10 groups of flies, Unpaired t-test). (b) The performance index for arabinose acuity was determined for starved *dArc^+/+^* (green) and *dArc^-/-^* (grey) flies using 100 mM arabinose in a place preference test (n ≥ 14 groups of flies, Unpaired t-test). (c) The performance index for sucrose acuity was determined for *dArc^+/+^* (green) and *dArc^-/-^* (grey) flies at different concentrations of sucrose in a place preference test (n ≥ 7 groups of flies, Two-way ANOVA). Subjects were mixed sex. Bar charts are presented as means ± s.e.m.

**Extended data figure 7.**
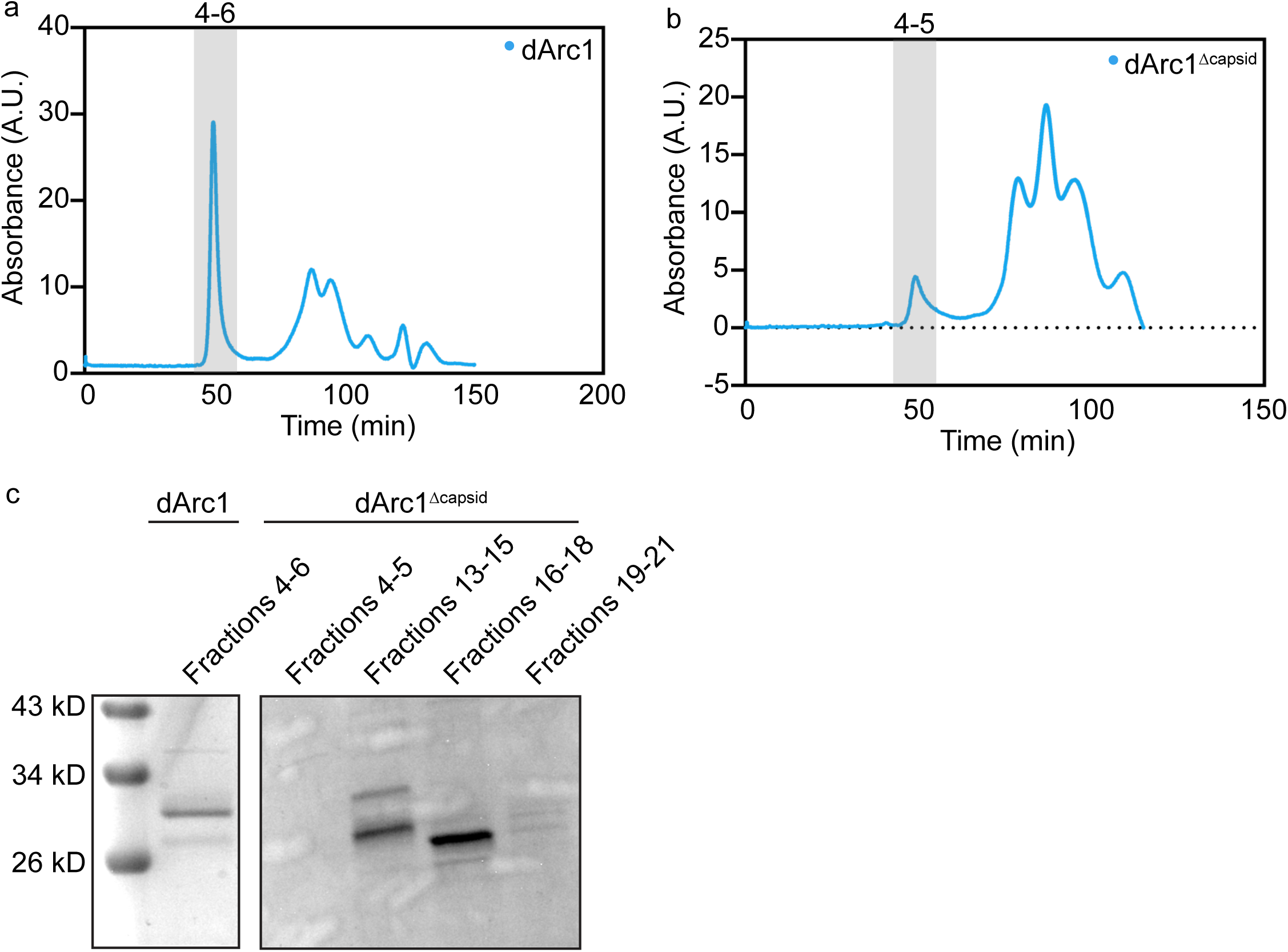
(a) Chromatogram of dArc1 purified protein showing the size exclusion chromatogram peaks selected for transmission electron microscopy. (b) Chromatogram of dArc1^Δcapsid^ purified protein showing the size exclusion chromatogram peaks selected for transmission electron microscopy. (c) Coomassie gel of the relevant peaks of purified dArc1 and dArc1^Δcapsid^ protein. Line charts are presented as a single purification.

**Extended data figure 8.**
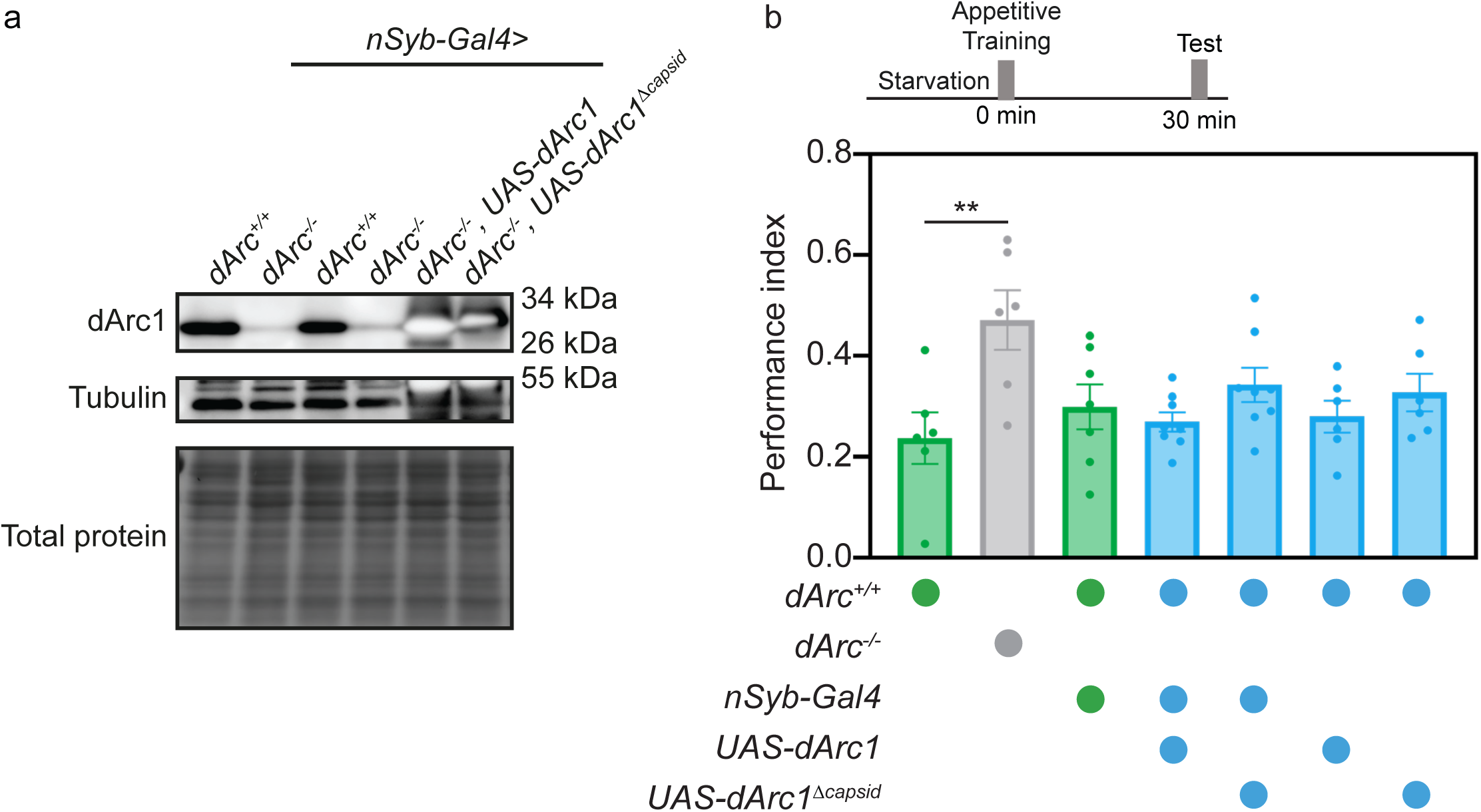
(a) Expression of dArc1 and dArc1^Δcapsid^ proteins in all neurons (nSyb-GAL4) of the fly in the *dArc^-/-^* background. (b) Starved *dArc^+/+^* flies expressing dArc1 and dArc1^Δcapsid^ proteins in all neurons (*nSyb-GAL4*) were subjected to an appetitive association paradigm using sucrose and tested after 30 minutes (n ≥ 6 groups of flies, One-way ANOVA). Subjects were mixed sex. Bar charts are presented as means ± s.e.m.

## EXTENDED DATA MATERIALS AND METHODS

### Quantitative poly chain reaction (qPCR)

Quantitative PCR (qPCR) was performed as previously described^1^. Briefly, total RNA was isolated from different samples using a standard Trizol (Qiagen) and chloroform extraction protocol, followed by ethanol precipitation. The precipitated RNA was treated with recombinant DNase I (Roche). Single-stranded cDNA was synthesized using the high-capacity cDNA reverse transcription kit (Applied Biosystems). Gene expression levels were determined using the PowerUp SYBR Green Master Mix for qPCR (Applied Biosystems) and measured with the Quantstudio 3 real-time PCR system (Applied Biosystems). Data normalization was performed using four housekeeping genes. The stability of these housekeeping genes was first assessed across samples, and the most stable genes were selected to normalize the expression of the target genes. The primers used for the RT-qPCR analysis are listed in the Extended Table.

### Western blot

Protein extracts were obtained using 1x RIPA buffer (50mM Tris-HCl (pH7.5), 1mM EDTA, 150mM NaCl, 0.1% SDS (sodium lauryl sulfate), 0.5% DOC (deoxycholic acid,sodium salt),1% Igepal CA-630 (NP40)). Protein extracts were mixed with 4X Laemmli buffer (40% glycerol, 250 mM Tris, 4% SDS, 50 mM DTT, pH 6.8) and heated at 95°C for five minutes. The proteins were separated by SDS-PAGE gel electrophoresis and transferred to a nitrocellulose membrane (GE Healthcare). Following transfer, membranes were briefly stained with Pierce™ Reversible Total Protein Stain and de-stained for total protein imaging. Membranes were blocked in 5% milk in 1X Tris-buffered saline-Tween (0.1%) (TBS; 10X: 152.3 mM Tris-HCl, 46.2 mM Tris-base, 1.5 M NaCl, pH 7.6) for 1 hour at room temperature. The blocked membranes were incubated with the primary antibody diluted in 5% milk in TBS-Tween overnight at 4°C. After incubation, membranes were washed five times for three minutes each with 1X TBS-Tween, followed by incubation with an HRP-conjugated secondary antibody (Jackson ImmunoResearch) in blocking buffer for one hour at room temperature. Membranes were then washed again three times for three minutes each time in 1X TBS-Tween, followed by two washes for three minutes in TBS. A chemiluminescence detection kit (Bio-Rad) was used to visualize the protein bands. Membranes were imaged using an Amersham ImageQuant 800 gel dock (Cytiva) and analyzed with ImageJ/Fiji (National Institutes of Health).

### Negative geotaxis assay

Negative geotaxis assays were performed following previously described protocols^1^. Flies were anesthetized 24 hours before the experiment to allow for clipping their wings. Flies were allowed to recover from anesthesia for at least one full day. Each experiment was conducted on a group of ten flies with clipped wings. During the assay, vials containing the flies were tapped on a hard surface to ensure that all the flies fell to the bottom of the vial. Negative geotaxis performance was calculated based on the percentage of flies that climbed seven centimeters within 15 seconds. Biological replicates (groups of 7-10 flies) of each genotype were tested across three technical replicates, and the average performance of each replicate was reported.

### Sensory acuity assays

Sensory acuity assays were performed on groups of 60 to 100 flies. In general, flies were selected prior to an experiment without sedation. For odor acuity assays, flies were given the choice to enter one of two tubes in a T-maze: one tube was scented with an odor (3-octanol or 4-methylcyclohexanol) while the other tube was unscented. Flies had two minutes to make a choice. For water acuity assays, flies were presented with a choice between a tube containing a piece of wet paper and another tube containing a piece of dry paper. Flies had two minutes to make a choice. For electric shock acuity assays, flies chose between two copper-lined tubes in a T-maze: one tube was electrified, and the other was not. Flies had two minutes to make a choice. For each assay, the number of flies in each tube was counted, and the performance index (PI) was calculated using the following formula:

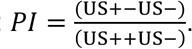

To determine the sugar acuity, flies were mouth pipetted into a dish divided into four quadrants: two quadrants were filled with a solution containing 1% agarose and the remaining two quadrants were filled with a solution containing 1% agarose and different concentrations of sucrose. Flies had five minutes to make a choice after which an image was taken. The number of flies were counted on the images, and the performance index (PI) was calculated based on the number of flies found in each quadrant and using the following formula:

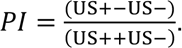

### Proboscis extension reflex

Flies were starved for 18 to 24 hours in a vial with a dampened Kimwipe. The proboscis extension reflex in response to a sucrose solution of was measured as described in^3^. Briefly, a fly is fixed in a 200 µl yellow tip with a slant cut to reveal the head. Wicks dipped in sucrose solutions were presented to the labellum of the fly until the fly extended its proboscis. Extension responses were scored manually under a stereomicroscope.

### Sensory neuron electrophysiology

Sucrose responses from the labellar taste sensilla were recorded using the extracellular tip recording method, as previously described^2^. Male flies aged 6 to 10 days were cold-anesthetized, and their proboscis was immobilized by inserting the reference electrode containing Beadle-Ephrussi Ringer solution through the thorax into the labellum. Neuronal firing rates of the L4 sensilla were captured using a glass electrode (10–20 μm diameter) filled with either 100 mM or 500 mM sucrose mixed with an electrolyte (30 mM tricholine). The glass recording electrode was connected to the TastePROBE system (Syntech) and a IDAC acquisition controller (Syntech). Signals were amplified (10x), band-pass-filtered (99–3000 Hz), and sampled at 12 kHz. To analyze neuronal firing rates in response to sucrose stimulation, spikes were manually sorted, and the number of spikes per stimulation was determined using the Autospike software.

### Sugar intake assay

Flies were first placed in a tube containing a dry piece of paper for 2 minutes. They were then transferred to a tube containing dried, blue-dyed sucrose paper for an additional 2 minutes. Immediately after exposure, flies were transferred to an Eppendorf tube and snap-frozen in liquid nitrogen. Flies were then visually scored for the presence of a blue proboscis or blue abdomen. Images of representative flies were captured using a Leica M165 FC microscope equipped with a Leica DMC 6200 camera and analyzed with the LAS X software.

### Triglyceride measurements

Total triglycerides normalized to total proteins were measured as described previously^4^. Briefly, two flies per biological replicate were homogenized in a lysis buffer (140 mM NaCl, 50 mM tris-HCl (pH 7.4), and 0.1% Triton X-100) containing a protease inhibitor cocktail (Thermo Fisher Scientific). The lysate was then used to determine protein and triglyceride concentrations. Protein concentrations were measured using the Pierce bicinchoninic acid (BCA) assay (Thermo Fisher Scientific; absorbance at 562 nm), while triglyceride concentrations were measured using the LiquiColor Test (Stanbio; absorbance at 562 nm).

